# DAPHNE: A global database of pest herbivores and their natural enemies

**DOI:** 10.64898/2026.09.28.754864

**Authors:** Daan Scheepens, Robin Freeman, Tim Newbold

**Affiliations:** Centre for Biodiversity and Environment Research, University College London, London, United Kingdom; Institute of Zoology, Zoological Society London, London, United Kingdom

## Abstract

Anticipating and managing the impact of herbivorous pests requires evidence-based knowledge about the interactions between pests and their natural enemies. Knowledge of species’ ecologies and biological interactions are typically disseminated in unstructured text across hundreds of thousands of scientific articles spanning a rapidly growing scientific literature, but manually screening this literature is time-consuming and impractical at scale. Taxonomic and geographic coverage thus remains sparse, and structured data on interactions between individual species are lacking, particularly for invertebrates. Large language models (LLMs) have recently emerged as a novel and powerful text-mining tool for knowledge extraction and evidence synthesis in a wide range of scientific disciplines, including ecology and conservation. In this study, we use an open-weight, pre-trained LLM to analyse abstracts of a corpus of published material on biological pest control to obtain a global Database of Pest Herbivores and their Natural Enemies (DAPHNE). This method resulted in the extraction of 175,359 species interactions from 112,830 published pest control studies, containing 16,881 unique animal taxa resolved at the species level. Interactions include herbivory (granivory, frugivory and gall-formation), predation, parasitism and hyperparasitism, and comprise a range of additional information, such as species’ taxonomy, pest status, pest importance, natural enemy importance, provision of biocontrol, associated plants and industries, invasiveness and the vectoring of pathogens and diseases. Validation of the results shows that most data columns were extracted with high recall and precision (*>*90%) and comparison of DAPHNE with other sources shows a high degree of agreement, but also highlights that the dataset contains information that may have been missed by other sources. In addition to the identification of important crop pests and their natural enemies, DAPHNE may be used for a variety of use-cases, including the identification of invasive species and pathogen vectors, trophic network analysis and literature search and discovery.

## 1 Introduction

Species interactions are a defining feature of ecological communities, underpinning population dynamics, community structure and ecosystem functioning (Paine, 1966). In agricultural landscapes, myriad interactions such as herbivory, disease-vectoring, predation and parasitism determine the delivery of agricultural services by natural enemies and disservices by herbivorous pests. Anticipating and managing these services and disservices requires reliable, evidence-based knowledge of these interactions.

Yet this knowledge remains difficult to assemble at scale. Predictions of ecosystem service and disservice delivery have traditionally relied on the manual classification of taxa by human experts (Karp et al., 2018; Martin et al., 2019; Millard et al., 2021; Oliver et al., 2015), but taxonomic and geographic coverage remains sparse (Cardoso et al., 2025), and interactions between individual species are lacking (Hortal et al., 2015; Poisot et al., 2021). Some resources may aid the identification of pests and their host crops (e.g., EPPO, 2026; García Morales et al., 2016) or interactions between species more broadly (Poelen et al., 2014), but knowledge of interactions remains sparse for many taxonomic groups. Furthermore, information on pest status, pest importance and the biological control provided by natural enemies is often dispersed across many different datasets or not in the public domain (e.g., CABI, 2026). This gap exists in large part because knowledge is typically disseminated in unstructured text in the scientific literature. Manually screening this literature is time-consuming and impractical at scale, which means that global syntheses of biological interactions have thus far been difficult to obtain.

Large language models (LLMs) have recently emerged as a powerful, novel text-mining tool for knowledge extraction and evidence synthesis in a wide range of scientific disciplines, including ecology and conservation science (Chang et al., 2025; Farrell et al., 2024; Mammides et al., 2025; Raeissi and Knapen, 2025). With careful design (Mammides and Papadopoulos, 2024; Moorthy et al., 2025), pre-trained LLMs have been shown to be capable of extracting and synthesising complex ecological information from text, including species identities, functional roles and geographical locations (Castro et al., 2024; Gougherty and Clipp, 2024; Scheepens et al., 2024), as well as species traits (Cornelius et al., 2025) and biological interactions (Keck et al., 2025). Moreover, LLMs have been used to classify documents in multilingual settings (Berdejo-Espinola et al., 2025a) and to retrieve evidence to answer specific conservation questions (Iyer et al., 2025). For many of these tasks, LLM performance has been found to match that of human experts.

In the domain of pest management, LLM-supported syntheses of species interactions, such as predator–prey or host–pathogen relations (Gougherty and Clipp, 2024; Scheepens et al., 2024), have the potential to aid in the natural control of crop pests and pathogens by providing farmers with evidence-based decision support (Rai et al., 2025; Xie et al., 2025). In addition, LLMs may help retrieve relevant evidence from queryable databases (Gallois et al., 2025; Iyer et al., 2025) to answer specific pest management questions. The benefit of synthesising any available evidence for any interaction is that individual pieces of evidence can be aggregated to obtain a rich and complex picture of a species’ ecology. By aggregating all the available evidence, an improved and evolving understanding of the ecology of species, and their ecosystem services and disservices, can be obtained.

In this study, we used the open-weight, pre-trained LLM GPT-OSS (OpenAI et al., 2025) to analyse the abstracts of 112,830 published, English-language studies on biological pest control to obtain a global Database of Pest Herbivores and their Natural Enemies (DAPHNE), which contains 175,359 individual interactions across 16,881 animal taxa resolved at the species level. These include herbivore–plant interactions (n=131,120), which comprise general herbivory (n=104,552), granivory (n=12,178), frugivory (n=9,649) and gall formation (n=4,741); and interactions between natural enemies and their prey or hosts (n=44,239), which comprise predation (n=18,095), parasitism (n=25,224) and hyperparasitism (n=810). Each interaction is further annotated with information such as the taxonomy of the species involved, pest status, pest and natural enemy importance, provision of biological control, associated plants and industries, invasiveness and pathogen vectoring. Together, this dataset provides a resource with relevance to a wide range of questions in pest management, invasive species biology, trophic network structure and community ecology.

## 2 Methods

### 2.1 Literature collection

Abstracts of published research articles on biological pest control were obtained from Scopus using the following search term: TITLE-ABS-KEY (“pest” OR “biological control” OR “natural enem*”) AND (LIMIT-TO (DOCTYPE, “ar”)) AND (LIMIT-TO (SUBJAREA, “AGRI”) OR LIMIT-TO (SUBJAREA, “ENVI”)) AND (LIMIT-TO (LANGUAGE, “English”)). This includes publications from agricultural, biological and environmental sciences, which includes, e.g., ecological studies, pesticide studies, bioassays, surveys of biocontrol agents and computer modelling studies of pest or biocontrol impacts. To reduce the computational cost of applying the LLM to every abstract, we used the R package gnfinder (Oldham, 2025, version 0.0.0.9000) to filter out abstracts that returned zero scientific names (e.g., Latin binomials) and thus likely do not contain any herbivores or natural enemies. It is possible, however, that abstracts containing only common names or substantial spelling or transcription errors may have been filtered out in this step. This yielded 112,830 abstracts potentially containing herbivores and natural enemies, spanning the years 1914 to 2025. Despite processing only English-language articles, the dataset exhibits decent global geographic coverage, although Africa is notably under-represented (Fig. 1).

### 2.2 Data extraction

To analyse the literature, we used the open-weight large language model GPT-OSS (OpenAI et al., 2025) with 120 billion parameters (i.e., neural network weights/connections), which was run locally on an Nvidia Run:ai server using the Python package ollama (version 0.6.1). This model, developed by OpenAI, is based on the Generative Pre-trained Transformer (GPT) architecture and has been shown to demonstrate strong reasoning performance across a wide range of different benchmarks (OpenAI et al., 2025). For each abstract in the literature collection, we first prompted the LLM to extract all herbivores and natural enemies (Latin binomials) from the abstract and article keywords, and then applied the LLM separately to each list of herbivores and natural enemies to extract further, species-specific information (Fig. 2). This two-step approach follows a least-to-most prompting strategy (Zhou et al., 2023) by dividing the complete extraction task into smaller, increasingly complex sub-tasks to improve model performance (Scheepens et al., 2024). We used GPT-OSS with its temperature parameter set at 1.0 (default) and its reasoning parameter set to ‘high’ to enable complex internal chain-of-thought (CoT) processing, and set the maximum generation token limit (num_predict; i.e., the maximum length of generated text) to -1 (unlimited) to allow the model’s reasoning architecture sufficient space to complete processing without premature truncation.

**Figure 1:**
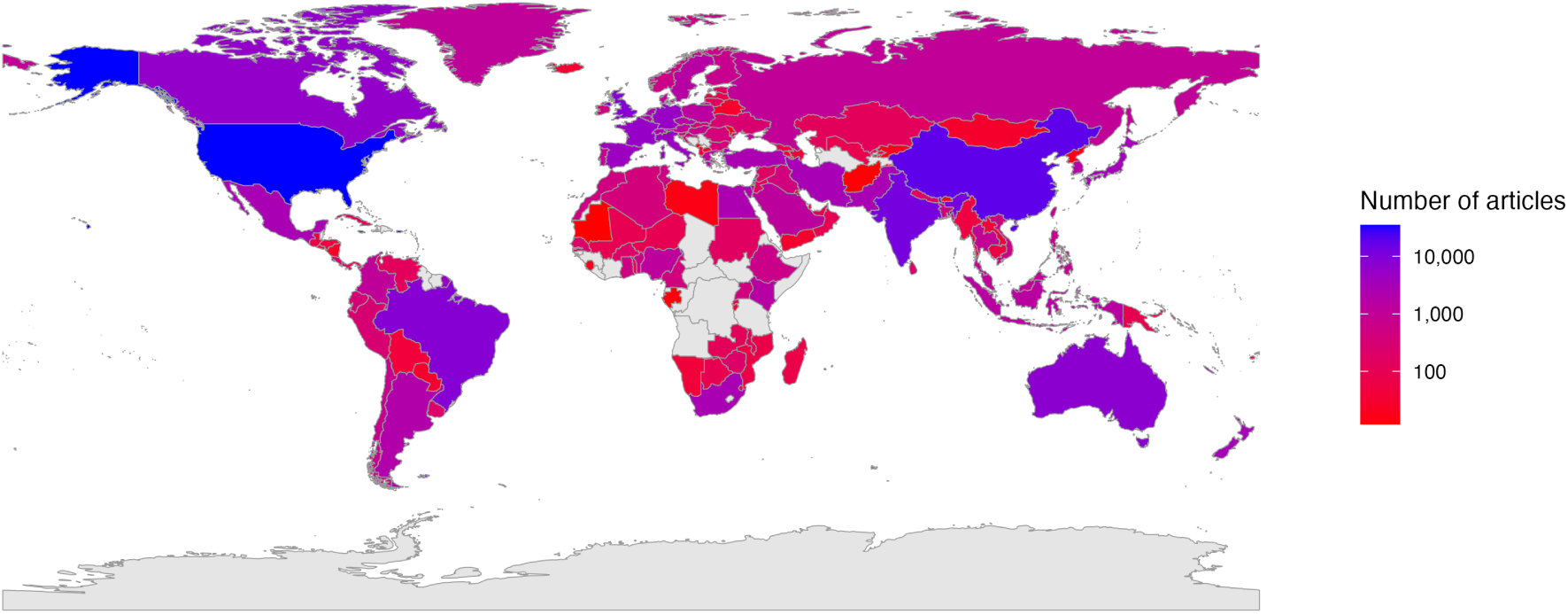
Geographic coverage of articles collected from SCOPUS using the search term: TITLE-ABS-KEY (“pest” OR “biological control” OR “natural enem*”) AND (LIMIT-TO (DOCTYPE, “ar”)) AND (LIMIT-TO (SUBJAREA, “AGRI”) OR LIMIT-TO (SUBJAREA, “ENVI”)) AND (LIMIT-TO (LANGUAGE, “English”)). The colour scale is plotted on a logarithmic scale.

**Figure 2:**
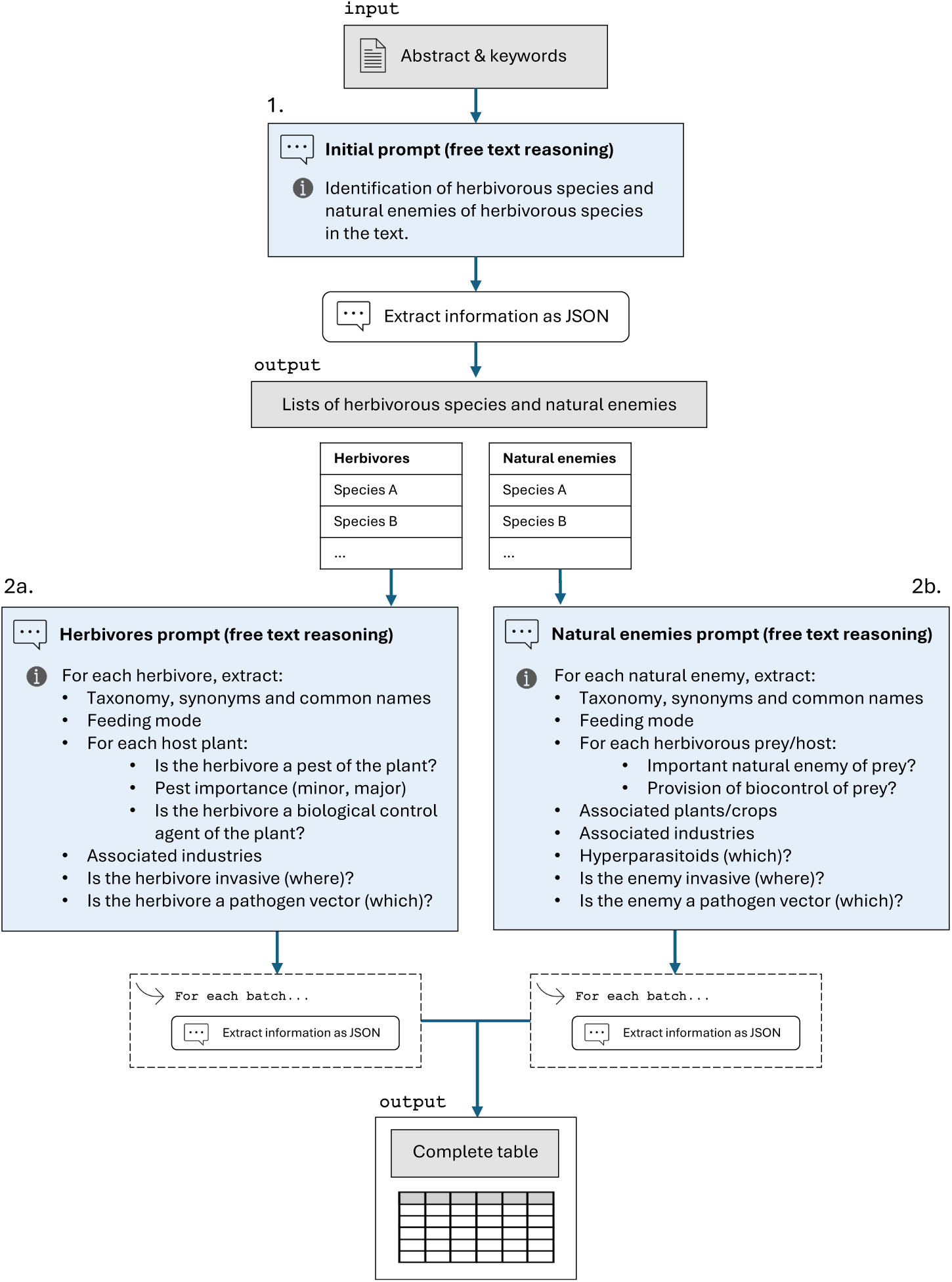
Simplified prompt schema used for the extraction of herbivorous pests and their natural enemies from the abstracts of published research articles. In the first step (1), the LLM was tasked with identifying all herbivores and natural enemies of herbivores in the text and returning these as lists in JSON format (for the full prompt see Fig. S1). In the second step (2), the LLM was provided the full prompt-and-response history, and either (a) the list of herbivores and a prompt specifying the further information to be extracted for herbivores, or (b) the list of natural enemies and a prompt specifying the further information to be extracted for natural enemies (for the full prompt see Fig. S2). Finally, the LLM was prompted to return the responses from step two in JSON format (in batches of five species), which were then merged into a complete table.

In both steps, we first prompted the LLM for a free-text response to allow the model to describe its reasoning, and then prompted the LLM again to return its generated response in a specified, machine-readable JSON format. Rather than generating the JSON table immediately, we found that the free-text reasoning step improved the ability of the LLM to respond accurately to complex questions, such as what suffices as evidence for predation, biological control, or whether a pest is stated to be of minor or major importance (Table 1). This reasoning step was saved in the database for full transparency and possible interrogation. For the free-text response, we used the following system prompt:

> “You are an expert ecologist and taxonomist and you are tasked with extracting information on herbivorous species and their natural enemies in scientific publications. You are to carefully follow the instructions given to you and are to extract information only from the text provided to you.”

**Table 1:** Information extracted per species, or specifically for herbivores (H) and natural enemies (NE). This information was extracted separately for herbivores and natural enemies using a multi-step prompting strategy (Fig. 2).

| Data Column | Description |
|---|---|
| Phylum | Taxonomic phylum of the species. |
| Class | Taxonomic class of the species. |
| Order | Taxonomic order of the species. |
| Family | Taxonomic family of the species. |
| Genus | Taxonomic genus of the species. |
| Species | Taxonomic species name (Latin binomial). |
| Synonyms | Synonyms associated with the species name (Latin binomial). |
| Common Name | E.g., the common name of Monecphora saccharina is the “sugarcane froghopper”. |
| (H) Feeding Mode | Whether the species is a herbivore through granivory (seed predation), frugivory (fruit consumption), gall-formation, or simply herbivory (de-fault; any other form of herbivory, or unspecified herbivory). |
| (H) Host Plants | List of plants or crops mentioned as host of the herbivore or as affected by it. |
| (H) Is Pest | For each host plant; whether the herbivore is a pest of this plant (True or False). |
| (H) Pest Importance | For each host plant; whether the pest is of minor or major importance. |
| (H) Is BCA | For each plant; whether the herbivore is stated to be a biological control agent of the plant. |
| (NE) Feeding Mode | Whether the species is a natural enemy through predation or parasitism (which includes parasitoidism). This may be inferred from the taxonomy (e.g., families of parasitoid wasps or predators). |
| (NE) Herbivore Prey | List of herbivorous species that are preyed upon or used as host by the natural enemy. |
| (NE) Important Enemy | For each prey; whether the natural enemy stated to be an important natural enemy of this prey/host (True or False). |
| (NE) Biocontrol | For each prey; whether the text explicitly states that the natural enemy provides biological control of this herbivorous prey/host (True or False). |
| (NE) Hyperparasitoids | List of hyperparasitoids that are stated to parasitise the natural enemy (if the natural enemy is a parasitoid). |
| (NE) Associated Plants | List of any plants associated with this natural enemy (e.g., where it is found to occur) or its herbivorous prey/hosts. |
| Invasive In | List of countries or regions in which the species is it stated to be invasive (if it is explicitly described with the term “invasive”). |
| Vectors | List of pathogens and/or diseases vectored by the species (if it is stated to be a vector of a pathogen or disease). |
| Associated Industries | List of any industries that are related to the host plants associated with the species and/or its herbivorous prey/hosts (if it is a natural enemy). |

Whilst many state-of-the-art LLMs are capable of returning responses in structured formats (Castro et al., 2024), GPT-OSS does not have an in-built functionality for structured output.

Instead, we prompted the model again to return its previous free-text response strictly according to a specified JSON schema. This was done in batches of five species at a time to prevent text truncation and ensure comprehensive outputs within the model’s generation token limits. For this step, the system prompt was changed to the following:

> “You are a JSON generator, and an expert ecologist and taxonomist. You are tasked with extracting information on herbivorous pests and their natural enemies in scientific publications. You are to carefully follow the instructions given to you and are to extract information only from the text provided to you. Return the information strictly according to the JSON format.”

### 2.3 Extracted information

The information extracted for each list of herbivores or natural enemies includes the species’ taxonomy insofar as stated in the abstract, any synonyms and common names, the feeding mode of the species with regards to their host (e.g., granivory, predation, parasitism) and a list of all host plants (in the case of herbivores) or herbivorous prey (in the case of natural enemies) explicitly stated in the text (Table 1). For each host plant of a herbivore, the LLM was then tasked with determining 1) whether the herbivore is stated to act as a pest of the plant, 2) whether the pest is of minor or major importance to this plant and 3) whether the herbivore is, instead, described as a biological control agent (BCA) of the plant (e.g., if the host plant is an invasive weed). Similarly, for each herbivorous prey or host of a natural enemy, the LLM was tasked with determining 1) whether the natural enemy is considered an important enemy, and 2) whether the enemy is explicitly stated to provide biological control, or has been successfully used a biocontrol agent, of the herbivore (for details, see Fig. S2).

The category ‘important natural enemy’ was devised to capture cases where a species was found to be an important enemy of a herbivore (e.g., a key parasitoid, a common or dominant enemy, a species responsible for the majority of predation in a study, etc.) even if it was not tested for pest regulation specifically, or not found to provide control of a pest but perhaps still plays a substantial regulative role within a larger guild of natural enemies. Biocontrol studies may posit that a natural enemy has “potential” to regulate a herbivorous pest, which we deemed to be insufficient evidence for biocontrol provision, but sufficient evidence for a species to be an important natural enemy. As such, all biocontrol agents are important enemies, but important enemies are not necessarily biocontrol agents. Natural enemies that were neither identified as ‘important’ nor as provisioning biocontrol were simply extracted as natural enemies of the herbivore.

Moreover, the LLM was tasked with identifying any hyperparasitoids parasitising the natural enemy, and what plants the natural enemy is associated with, if any were stated in the text. Finally, for both herbivores and natural enemies, the prompt instructed the LLM to return a list of industries associated with the species or its host plants (e.g., arable crops, greenhouse production, pasture, forestry and timber production) and determine whether the species is invasive (and if so, where?) and whether the species vectors any pathogens or diseases (and if so, which?). Complete descriptions, as embedded in the JSON schemas, can be found in the supplementary information (Table S1).

My prompts did not distinguish between potentially different qualities of evidence for a piece of extracted information. This means that the LLM was instructed to extract a piece of information, regardless of whether this information was experimentally verified in the abstract (e.g., “our experiments showed that species X regulated species Y”), or asserted without experimental evidence (e.g., “species X is widely used as a biocontrol agent of species Y”). Distinguishing between such types of evidence can become highly ambiguous and was deemed beyond the scope of this study. However, since the corpus used in this study comprises peer-reviewed research publications, asserted statements are expected to reflect scientific consensus.

### 2.4 Prompt fine-tuning and model evaluation

To validate the results, we manually checked the output generated by GPT-OSS against two manually labelled datasets: a training set and a held-out test set, both comprised of 100 randomly selected abstracts. These sets contained 873 interactions between herbivores and plants, and between natural enemies and herbivores (628 in the training set and 245 in the test set). The training set was used during the creation of the prompts to identify errors and subsequently fine-tune the prompts to improve the generated results. Effective prompting is highly model-dependent, and usually a process of trial-and-error (Cao et al., 2024; Razavi et al., 2025; Wang et al., 2024). After a series of fine-tuning cycles, the prompts assumed the structure as shown in Fig. S1 and Fig. S2. Two instructions that the model consistently performed poorly on were related to the data columns ‘Important Natural Enemy’ and ‘Biological Control’ for natural enemies. This was related to difficulty in distinguishing between species that were stated to have “potential” for biocontrol (or candidates for biocontrol measures) and species that were experimentally verified to effectively regulate a pest or to be successfully utilised as a biological control agent. Only the latter was deemed sufficient evidence for biological control, whilst the former was characterised as evidence for the species being an important natural enemy (which may reflect a potential for pest regulation). Furthermore, the model would struggle to correctly reason that if one species was found to be an important natural enemy, and a group of other species were found to be equally effective predators, then it follows that these species should also be identified as important natural enemies. By prompting GPT-OSS to improve the natural enemies prompt (prompt 2b in Fig. S2) with regards to these instructions, the model added a short “cheat-sheet” to the prompt, which was found to improve its ability to generate the correct results.

We used four metrics to evaluate model performance: proportion correct (PC; also referred to simply as accuracy), precision, recall and F1-score (Eq. 1). Whilst PC measures overall predictive performance as the number of true predictions relative to the total number of predictions, precision measures the ability of the model to correctly predict a true positive (and is thus penalised by the number of false positives), recall measures the ability of the model to correctly capture all true positives in the dataset (and is thus penalised by the number of false negatives), and F1-score is the harmonic mean of precision and recall. For string variables (e.g., species taxonomy; see Table S1), positives refer to cases where a string is non-empty or non-NA and negatives refer to cases where a string is empty or NA. For boolean or categorical variables, we treated the evaluation as a multi-class classification problem with ‘None’ as an extra class to designate empty or NA labels: for these variables, model performance metrics are reported as weighted averages across all classes (e.g., True, False and None for boolean variables).

For taxonomic information, minor differences between predictions and manual labels may not be indicative of actual errors, but may instead be a result of misspellings in the text that were extracted verbatim by the LLM but were corrected in the manual labels, or a correct family returned as order, etc. To differentiate such cases, we manually identified mismatches in the training and test set as either minor or major, where minor mismatches indicate differences in spellings, shifts in the correct taxonomy (e.g., the correct class extracted as phylum, etc.) or missing information in the text that was correctly inferred by the model (e.g., the inference of *“Diuraphis noxia”* rather than *“D. noxia”* as stated verbatim in the text), and major mismatches refer to anything else. We then computed proportion-correct once as PC_exact_, which counts all mismatches as errors, and once as PC_approx_, which only counts major mismatches as errors. In addition, we manually differentiated cases in which the taxonomic information was available in the text 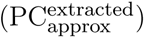 versus cases in which the taxonomy was not available in the text and thus had to be inferred by the LLM from its training data 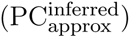, which we assume to be more error-prone and not strictly indicative of named-entity recognition capability.

### 2.5 Post-processing

The extracted data were post-processed to correct taxon binomials and harmonise names and taxonomies with the Global Biodiversity Information Facility (GBIF: The Global Biodiversity Information Facility, 2026). In this way, many taxonomic ranks could be filled in where they were not extracted by the LLM (e.g., missing from the text) and/or corrected where the extracted term was incorrect or obsolete. These corrections were made to herbivores and natural enemies, but not host plants as this column contains many cases of common names or non-specific terms (e.g., “greenhouse vegetables”). Instead, users are recommended to string-match host plants with a series of terms rather than single Latin binomials to maximise matches (e.g., maize, corn, zea mays; see Table S3). Taxa were first cleaned to remove any suffixes such as “sp.”, “nov.”, “indet.”, etc., and were then matched with accepted names on GBIF. In cases of unmatched binomials, we manually checked the respective abstracts and the LLM reasoning output for errors (e.g., the wrong combination of genus and species), and searched online resources for possible synonyms or misspellings (e.g., EPPO, 2026; García Morales et al., 2016; GBIF: The Global Biodiversity Information Facility, 2026; VanDyk et al., 2026). The removal of the suffix “sp. nr.” (species near) means that evidence related to a taxon “near” a particular species is aggregated into the evidence for this particular species, as they are assumed to be highly similar or closely related. For transparency, however, the original names as stated verbatim in the abstract were retained in the database.

### 2.6 Summarising evidence

The information extracted in DAPHNE may be ranked in terms of certainty or reliability through the number of abstracts (identified by a unique electronic identifier; EID) providing evidence for an interaction (e.g., herbivory of species *X* or host plant *Y*) or any additional information (e.g., a pest to be of major importance). For example, an interaction extracted from 100 abstracts may be assumed to be more reliable than an interaction extracted from a single abstract. A single extraction of an interaction may still be informative, since low representation may also be suggestive of research bias, but the information is more likely to be erroneous (and should be manually checked). Notwithstanding research bias, a higher number of abstracts evidencing an interaction may also be indicative of ecological, economic or agricultural importance.

We summarised the evidence for all herbivore–plant and enemy–herbivore interactions by summing the number of abstracts evidencing the interaction (nEID). Evidence for natural enemies was then further summed for important enemies (nImportantEnemy). Rather than summarising evidence between herbivores and all individual plants (which are referenced with many different names in the database, and often as broader groups rather than single taxa), we summarised evidence for 77 crop groups as defined by the Food and Agriculture Organization (FAO Food and Agriculture Organization (FAO), 2005), which include cereals, vegetables, fruits and nuts, oilseeds, roots and tubers, beverage and spices, legumes, sugar, and others. We have also included five widespread tree species used in plantation forests: pine (Pinus spp.), eucalyptus (Eucalyptus spp.), douglas-fir (Pseudotsuga menziesii), spruce (Picea spp.), and teak (Tectona grandis). These crops and tree species were then string-matched using their common names and Latin binomials (Table S3). For herbivores, we then further summed the number of abstracts evidencing the herbivore to be a pest (nPest) and a major pest (nMajor).

We then examined the agreement of DAPHNE with pest control classifications from Oliver et al. (2015), classifications of predators, parasitoids, herbivores and herbivore pests and non-pests from Martin et al. (2019), and pests of 79 of the 82 string-matched crop and tree species as listed in the EPPO database (EPPO, 2026) (no EPPO host lists were available for the remaining three crop groups: mixed cereals, sweet sorghum and grass). The species from these sources were matched with species in DAPHNE to test the extent to which species classified in Oliver et al. (2015) as providing pest control as a primary function are represented in the database with at least one evidence of acting as an important natural enemy of a pest (nImportantEnemy *>* 0); and species classified as predators, parasitoids and herbivores in Martin et al. (2019) were extracted with these same labels in DAPHNE, and whether pest herbivores are associated with at least one evidence of major pest importance (nMajor *>* 0) and non-pest herbivores are not associated with major pest importance (nMajor = 0). Lastly, we tested the extent to which species listed in the EPPO database as host or major host of a particular crop or tree species were captured in DAPHNE as pests of this plant (nPest *>* 0 or nMajor *>* 0). Agreement was then computed from those source taxa contained in DAPHNE as the percentage of taxa for which the respective classification matched. To assess whether filtering DAPHNE to exclude interactions with low evidence counts results in better agreement, we computed the agreement between DAPHNE and the external sources for five different selection criteria; nEID *≥* 1 (no filtering), and nEID *≥* 3, 5, 10 and 20.

The pest control classifications in Oliver et al. (2015) were not compared against the DAPHNE ‘Biocontrol’ field, as the definition of biocontrol used in this study (demonstrated control or successful use as a biocontrol agent) is substantially stronger than the definition used in Oliver et al. (2015), where it is defined more broadly as “likely to act as natural enemies of crop pests” – which aligns better with DAPHNE’s ‘ImportantEnemy’ category. The classifications in Martin et al. (2019) were grouped according to the following groups: pest herbivores (“pest herbivore”, “larval pest herbivore, adult pollinator (adults)”, “larval pest herbivore, adult pollinator”); non-pest herbivores (“non-pest herbivore”, “larval non-pest herbivore, adult pollinator”); predators (“predator”, “aphid-tender, predator”, “larval predator, adult pollinator”); and parasitoids (“parasitoid”, “parasitoid of bees”, “larval parasitoid, adult pollinator”).

## 3 Results

Across 112,830 abstracts of published research articles, our method extracted 175,359 total interactions. Of these, 131,120 are interactions of herbivory between herbivores and plants (104,552 general or undefined herbivory, 9649 frugivory, 12,178 granivory and 4741 gall-formation interactions), 43,429 are predatory interactions between natural enemies and herbivores (18,095 predation; 25,224 parasitism; and 110 with feeding mode unknown/NA), and 810 are interactions of hyperparasitism between hyperparasitoids and parasitoid natural enemies (Figure 3). In total, 16,881 unique taxa were identified and resolved at the species level, of which 15,089 could be harmonised with GBIF. Of these, 759 species were identified as invasive in a particular country or region (not necessarily the location of the study), and 653 species were identified as vectoring a pathogen or disease.

**Figure 3:**
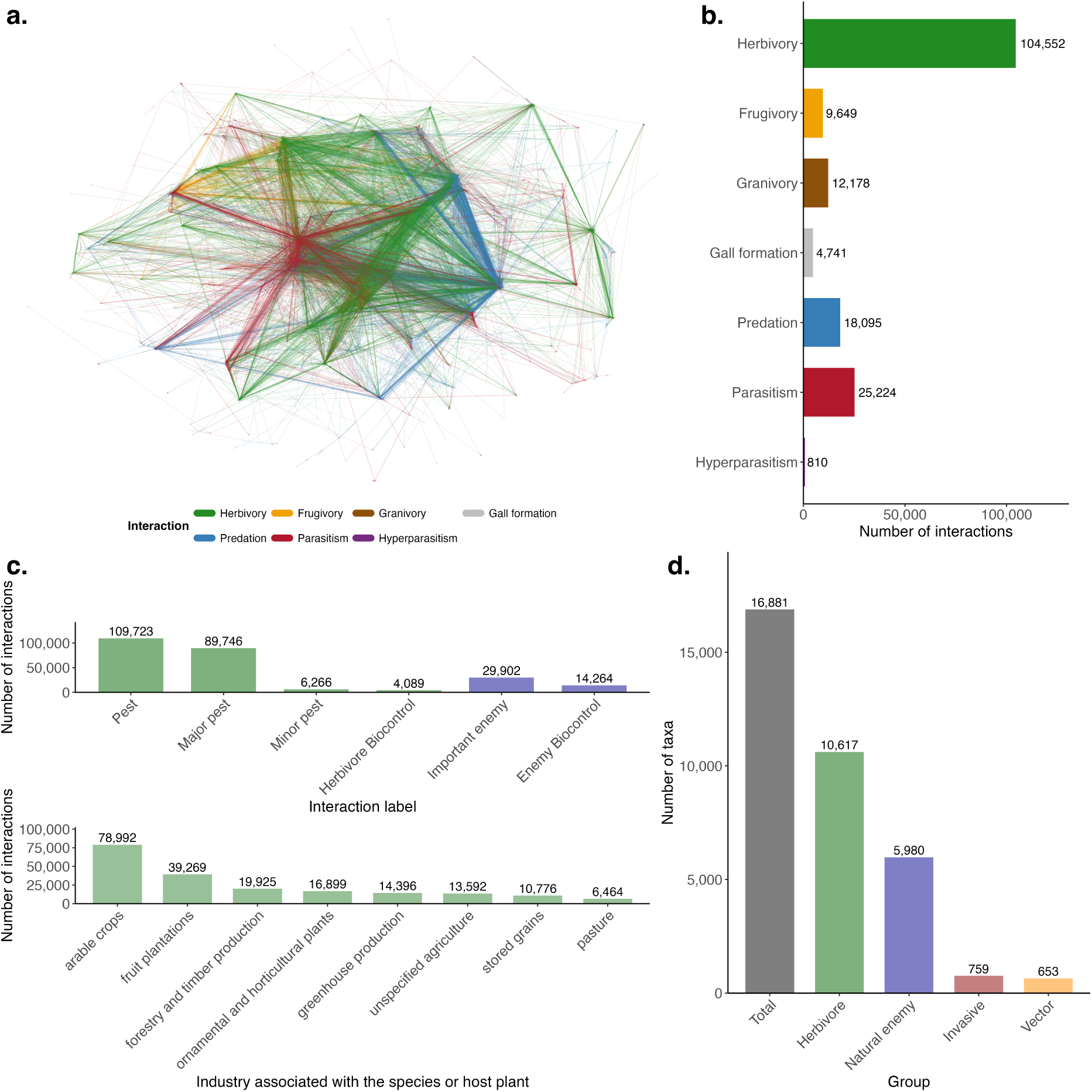
General overview of the DAPHNE database. Panel a: visualisation of the interaction network spanned by all interaction types in DAPHNE (only the largest connected component is shown here). Panel b: the number of interactions for each interaction type. Panel c: the number of interactions annotated with additional herbivore-specific information (pest, major pest, minor pest or herbivorous biocontrol agent) or additional enemy-specific information (important enemy and biocontrol provision). Panel d: number of unique taxa (total, or classified as herbivores, natural enemies, invasive and disease-vector).

The majority of extracted species are insects, although arachnids, mammals and birds comprise a substantial proportion of predators (Fig. 4). Arachnids and mammals are also represented to a larger degree among herbivores, as are nematodes (e.g., phytophagous root-knot nematodes) and molluscs. Nematodes are also represented among parasites, as well as a small number of arachnids.

**Figure 4:**
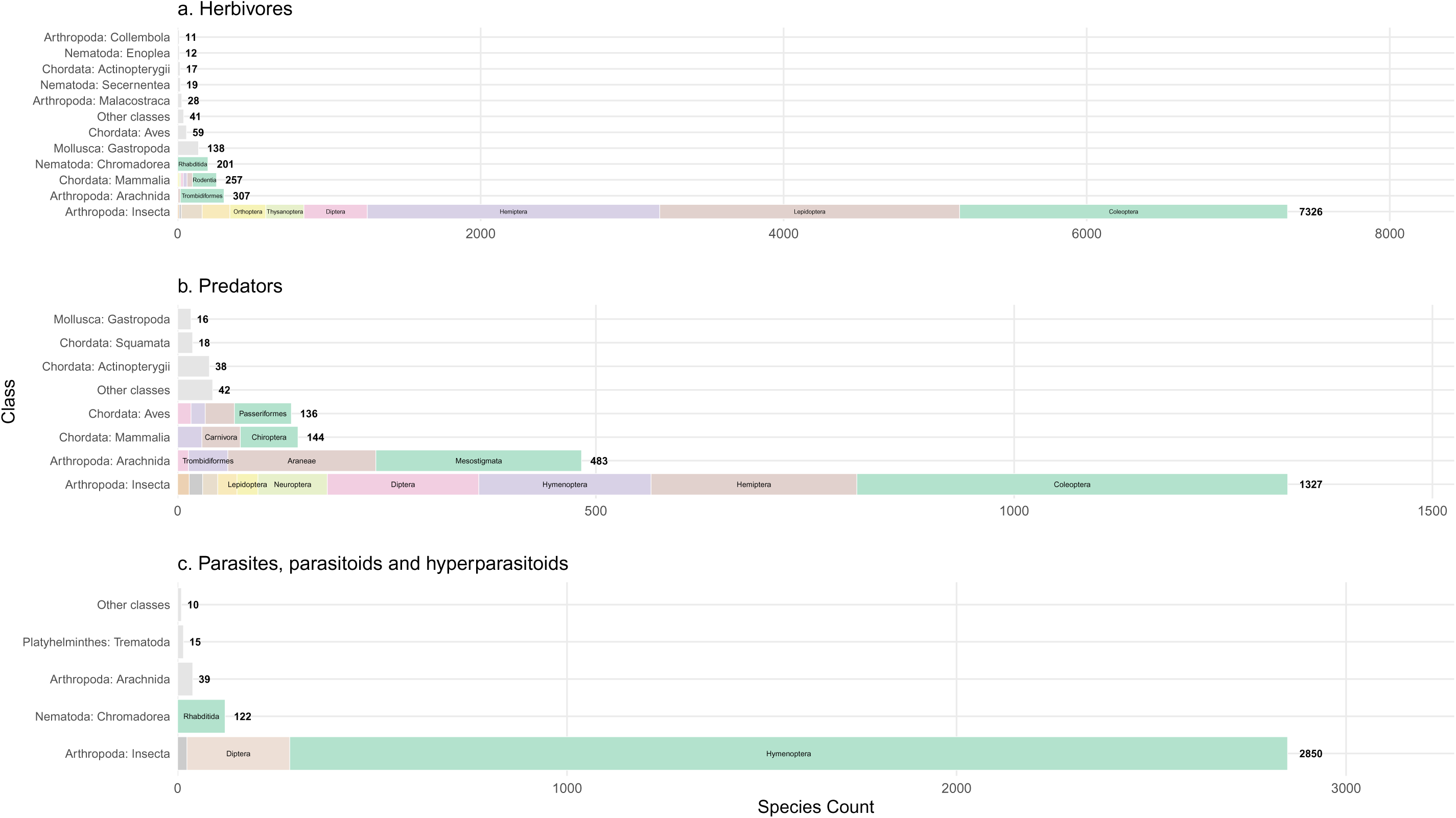
The number of unique species extracted per taxonomic class (following binomial corrections and harmonisation with GBIF), shown for herbivores (a), predator (b) and parasites, parasitoids and hyperparasitoids (c). Classes with fewer than 10 species counts were aggregated into ‘Other classes’. The figure shows that the majority of extracted species are insects, followed by arachnids (herbivores and predators) and mammals and birds (predominately predators; largely comprised of bats and passerines).

### 3.1 Model validation

The performance of GPT-OSS on this task was high on most data columns (Table 2) and mostly showed only minor variation between the training and test set (Table S2). Indeed, in many cases performance on the test set exceeded performance on the training set, suggesting that the prompt fine-tuning did not ‘overfit’ to the training data.

**Table 2:** Accuracy obtained by the model on each data column, on all manually labelled abstracts from the training and test sets. Due to a low number of positive labels in some of the data columns in the training and test set, the results are shown here for both sets combined, but separate evaluations are presented in Table S2. Taxonomic information was evaluated as proportion-correct (PC; Eq. 1), using either exact matches (PC_exact_) or approximate matches (PC_approx_). Approximate matches were further computed for taxonomic terms that were directly extracted from the text 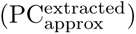 and for terms that were not available in the text and thus had to be inferred by the model 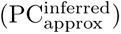. All other columns were evaluated with PC, precision (Pre), recall (Rec) and F1-scores (Eq. 1). In the case of boolean and categorical variables, these metrics are reported as weighted averages across all classes. The support for each each score (*N*) designates the total number of true positives, false positives, true negatives and false negatives. Herbivore-specific data columns are denoted with (H) and natural-enemy-specific columns are denoted with (NE).

| Data Column | PC<sub>exact</sub> | PC<sub>approx</sub> | PC<sup>extracted</sup><sub>approx</sub> | PC<sup>inferred</sup><sub>approx</sub> | *N* |
|---|---|---|---|---|---|
| Phylum | 95.5 | 95.5 | 100 | 94.9 | 804 |
| Class | 98.6 | 98.6 | 100 | 97.8 | 804 |
| Order | 99.5 | 99.5 | 100 | 98.4 | 804 |
| Family | 96.6 | 97.8 | 99.7 | 91.3 | 804 |
| Genus | 99.3 | 99.8 | 99.8 | - | 804 |
| Species | 98.6 | 99.1 | 99.1 | - | 804 |
| Data Column | PC | Pre | Rec | F1 | *N* |
| Synonyms | 99.3 | 64.7 | 100 | 78.6 | 804 |
| Common Name | 86.1 | 59.9 | 100 | 74.9 | 804 |
| Species Type | 99.9 | 99.9 | 99.9 | 99.9 | 804 |
| Feeding Mode | 99.5 | 99.6 | 99.5 | 99.5 | 804 |
| (H) Host Plants | 95.9 | 97.7 | 97.7 | 97.7 | 392 |
| (H) Is Pest | 93.1 | 93.3 | 93.1 | 93.2 | 392 |
| (H) Pest Importance | 92.9 | 93.1 | 92.9 | 92.9 | 392 |
| (H) Is BCA | 95.7 | 95.7 | 95.7 | 95.7 | 392 |
| (NE) Herbivore Prey | 97.6 | 98.6 | 98.6 | 98.6 | 657 |
| (NE) Important Enemy | 92.5 | 92.5 | 92.5 | 92.5 | 657 |
| (NE) Biocontrol | 94.4 | 95.5 | 94.4 | 94.7 | 657 |
| (NE) Hyperparasitoids | 100 | 100 | 100 | 100 | 548 |
| (NE) Associated Plants | 98.6 | 98.2 | 100 | 99.1 | 567 |
| Invasive In | 99.8 | 90.0 | 100 | 94.7 | 811 |
| Vectors | 99.8 | 100 | 83.3 | 90.9 | 804 |
| Associated Industries | 88.9 | 90.8 | 88.8 | 89.4 | 891 |

Species types (herbivores vs. natural enemies) were accurately distinguished, with only a single mistake in the training set (S6; related to a gall wasp mistakenly identified as a natural enemy). The types of interactions (‘Feeding Mode’ labels) were also extracted with excellent accuracy, precision and recall scores in both the training and the test set: the LLM was able to extract predation and parasitism perfectly, but mislabelled four out of 288 cases of herbivory (Fig. S7a). Pest-specific information was extracted with high levels of accuracy, with LLM extractions obtaining host plants with 97.7% precision, pest labels with 93.3% precision, pest importance labels with 93.1% precision and biological control agent labels with 95.7% precision across the training and test set (Table 2). True labels of pests were extracted with a slightly better precision than False labels (96.3% vs. 91.6%; Fig. S8a) and major importance labels were extracted with a better precision than minor importance labels (91.6% vs. 73.7%; Fig. S9a), whilst precision was comparable for True and False labels of herbivorous biological control agents (100% vs. 97.3%; Fig. S10a).

For natural enemies, LLM extractions obtained herbivore prey species with 98.6% precision, important natural enemy labels with 92.5% precision, and biocontrol provision labels with 95.5% precision across both the training and test set (Table 2). With respect to true labels only, important natural enemy extractions were made with a considerably higher precision (93.3%; Fig. S11a) than the extractions of biocontrol provision (65.7%; Fig. S12a), which was comparable across the training and the test set (Fig. S11 and Fig. S12). False biocontrol provision labels, however, were extracted with a considerably higher precision of 99.0%, and are thus likely to be more reliable across the whole dataset. Furthermore, all 96 instances of hyperparasitism in the training and test set were correctly extracted (100% precision and recall) and plants associated with the natural enemies were correctly identified with a precision of 98.2% and recall of 100% (Table 2).

All cases of invasiveness (region or country where a species is stated to be invasive) were identified correctly (100% recall) but there was one case where the phrase “two invasive pests of tomato and canola in Europe and North America, respectively” was interpreted by the LLM to mean that both pests were invasive in Europe and North America, whilst only one is invasive in Europe and the other in North America (reducing the precision to 90%). All pathogen- or disease-vectors extracted by the model were correct (100% precision), but the model missed two vectors of grapevine leafroll disease related to a single abstract (83.3% recall). The performance with which industries associated with the herbivore or natural enemy were extracted was highly variable between industry-types. Whilst most industries were extracted with high precision, the model struggled with the identification of pasture (65.5% precision; Fig. S13a).

Based on approximate matches, taxonomy was extracted by the model with accuracy of at least 99.1%, and, where it was missing in the text, was inferred with an accuracy of at least 91.3%. Synonyms and common names were predicted with relatively low precision scores of 64.7% and 59.9%, respectively. In the case of common names, this is primarily the result of a large number of LLM extractions of generic common names (e.g., ground beetle, mirid bug), which were not extracted as manual labels. There were only 11 manually extracted cases of synonyms as stated in the text, all of which the LLM correctly identified. At the same time, however, the LLM erroneously extracted six additional synonyms from an abstract in which the authors describe six species that were previously misidentified as other species (the misidentifications of which the LLM erroneously extracted as synonyms).

### 3.2 Comparison with other sources

#### 3.2.1 Pest control

Out of 160 species classified in Oliver et al. (2015) as likely to exert pest control, 140 species (87.5%) obtained at least one count of evidence for acting as an important natural enemy of a herbivore (nImportantEnemy *>* 0). It should be noted, however, that absence of evidence in DAPHNE should not be interpreted as evidence for absence; indeed, of the 20 natural enemies that obtained no ‘ImportantEnemy’ evidence in DAPHNE, most species (13) were associated with a single piece of evidence in the database (with a mean EID count of 1.95), and it is thus likely that additional evidence may attest to their importance as natural enemies of pest herbivores. For this reason, a higher agreement between Oliver et al. (2015) and DAPHNE is observed when the database is filtered for nEID *≥* 3 and higher (Fig. 5), although the number of compared taxa decreases rapidly.

**Figure 5:**
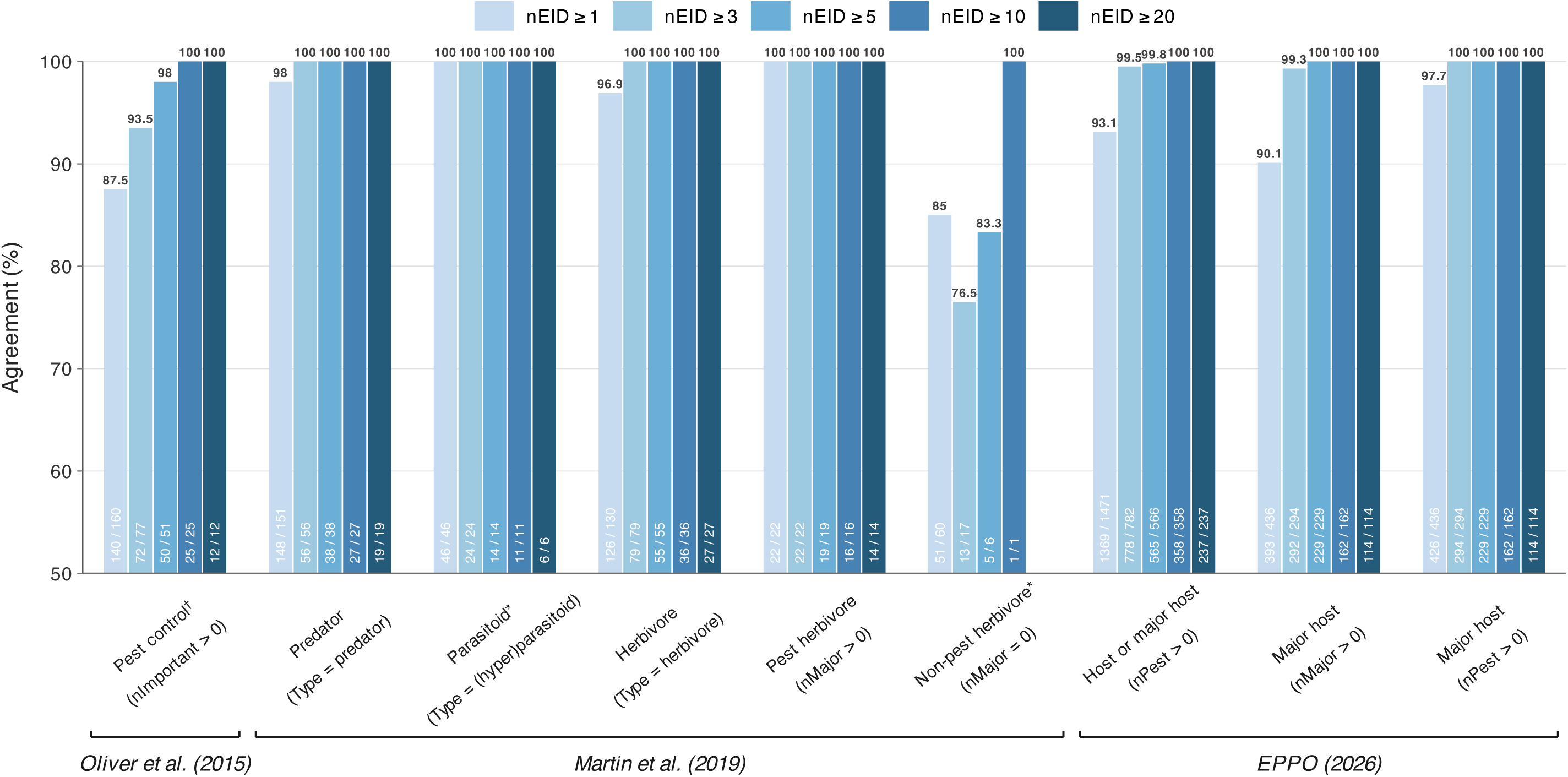
The % agreement between DAPHNE and three other data sources (EPPO, 2026; Martin et al., 2019; Oliver et al., 2015) for 9 classifications of species or their interactions, computed as the number of taxa that could be matched in DAPHNE for which the source classification matched with the respective DAPHNE data summary (shown in brackets). The figure shows the % agreement for five different filters on the DAPHNE database, which were applied prior to computing agreement; namely, the number of individual publications (nEID) evidencing an interaction to be greater or equal to 1 (i.e., no filter), 3, 5, 10 and 20. Disagreement does not necessarily suggest that the DAPHNE extractions are erroneous, however, and for many mismatches evidence was found to support the data entry in DAPHNE. Some of these mismatches were found to result in an unfair comparison, and were thus removed from the category (marked with an asterisk). Nevertheless, some data entries in DAPHNE were found to be either erroneous, exaggerated or the result of highly ambiguous language, but the figure shows that such cases are rapidly filtered out by filtering the database to exclude interactions with low evidence counts (e.g., using the criterium nEID *≥* 3). *^†^*pest control in Oliver et al. (2015) was defined as species that are likely to act as natural enemies of crop pests, and was thus compared to the DAPHNE ‘ImportantEnemy’ variable rather than the ‘Biocontrol’ variable, which was given a much stricter definition of demonstrated control or utilisation as a biocontrol agent.

#### 3.2.2 Predators, parasitoids and herbivores

Out of 151 species classified as predators in Martin et al. (2019), 148 were extracted in DAPHNE as natural enemies with feeding mode ‘predation’ (98.0%); two predators were extracted exclusively as herbivores in DAPHNE, and one species was exclusively extracted as a parasitoid. No evidence for predation could be found in other online sources for the two herbivores (*Kleidocerys resedae* and *Subcoccinella vigintiquatuorpunctata*), which suggests that the DAPHNE extractions are accurate. The parasitoid mismatch refers to the rove beetle *Aleochara sparsa*, which is mentioned in the literature as a larval ectoparasitoid of carrot fly, *Psila rosae*, but are predatory as adults. Evidently, the model extracted evidence for the former but not the latter. All three species were poorly evidenced in the database (nEID = 1), and were thus filtered out with a stricter selection criterium (nEID *≥* 3), which then resulted in 100% agreement between the datasets (Fig. 5).

Parasitoids in Martin et al. (2019) were classified as any natural enemy parasitic on other arthropods at any life stage, and thus may include hyperparasitoids. Out of 48 species classified as parasitoids, 43 species were extracted in DAPHNE as natural enemies with feeding mode ‘parasitism’ (one of which was also extracted with feeding mode ‘predation’) and 3 with feeding mode ‘hyperparasitism’. Two species (*Oscinella frit* and *Trogoderma glabrum*), which were classified in Martin et al. (2019) as parasitoids of bees, were extracted in DAPHNE as herbivores. We could not find any evidence of these species acting as parasitoids, the former being a major agricultural pest and the latter, although identified in one external study as associated with mason bee nests, but not as an important parasite (Zajdel et al., 2014), is predominantly a pest of stored products. Based on this assessment, we removed these two species from the comparison, which resulted in an agreement of 100% between the two databases (Fig. 5).

From 130 species classified as herbivores in Martin et al. (2019), 126 were extracted as herbivores in DAPHNE (96.9%), whilst 4 species were extracted exclusively as natural enemies. These species (*Amara bifrons*, *Amara familiaris*, *Amara ovata* and *Harpalus distinguendus*) are predominantly granivorous but the literature confirms that they may predate opportunistically. As their roles as natural enemies were evidenced in each case by a single abstract (nEID = 1), it is likely that their more important roles as herbivores would be evidenced by additional literature. Higher selection criteria (nEID *≥* 3 and higher) thus rapidly filtered these mismatches out (Fig. 5).

#### 3.2.3 Pest and non-pest herbivores

Out of 22 species classified as pest herbivores in Martin et al. (2019), all 22 (100%) obtained at least one count of major pest evidence (nMajor *>* 0). Conversely, 108 species classified as non-pest herbivores herbivores could be matched with taxa in DAPHNE, but more than half of these (57 species) obtained major pest evidence (nMajor *>* 0). However, these mismatches were found to include some globally important pests such as *Tribolium castaneum* (red flour beetle), *Apolygus lucorum* (green plant bug) *Hypera postica* (alfalfa beetle), *Phyllotreta cruciferae* (crucifer flea beetle), *Sitona lineatus* (pea leaf weevil) and others. Manual verification of the associated abstracts validated that the evidence indicated pest status for 48 out of 57 species in at least one instance. The remaining 9 non-pest herbivores obtained only one (six species) or three (three species) instances of major pest evidence in DAPHNE, which manual verification showed to be weak or exaggerated, and which may thus be considered erroneous extractions.

It is likely that the mismatch for the 48 pest species resulted from the fact that Martin et al. (2019) looked at pests in crop fields, whilst DAPHNE included other industries such as forestry. The comparison is, furthermore, made difficult by the fact that species may be pests in one region (e.g., where they are non-native) but not in another; our methodology extracted species as pests in myriad countries and regions across the world, whereas the Martin et al. (2019) study was conducted in Europe. These 48 species were thus considered to result in an unfair comparison, and were removed from the non-pest herbivore category. Due to the fact that the remaining non-pest herbivores obtained very low evidence counts in DAPHNE (average nEID of 2.2, median of 1), the number of compared taxa decreased rapidly with increasingly strong selection criteria (Fig. 5). This suggests, however, that filtering out interactions with low evidence counts should be expected to filter out non-pest herbivores much more rapidly than pest herbivores (which are, naturally, studied much more extensively in the biological pest control literature).

#### 3.2.4 Major pests

Across the 79 host crops and tree species with EPPO host lists that were pattern-matched with host plants in DAPHNE, 93.1% of species listed as either “host” or “major host” of a crop in the EPPO database (EPPO, 2026) obtained at least one count of pest evidence (nPest*>*0) in DAPHNE (Table S4). For major hosts specifically, 97.7% obtained at least one count of pest evidence (nPest*>*0), and 90.1% obtained at least one count of major pest evidence in DAPHNE (nMajor*>*0). Overall, species labelled as major host in EPPO tended to obtain marginally higher nPest counts (mean=62.0) and nEID counts (mean=66.7) than hosts (59.4 and 64.3, respectively), which suggests that increased evidence counts may be indicative of increased pest importance, although this was highly variable between crops (Table S4) and assumes the EPPO distinction between hosts and major hosts to be accurate. Mismatches were mostly restricted to interactions with low evidence counts, which means that stricter selection criteria (nEID *≥* 3) resulted in much higher agreement (Fig. 5).

## 4 Discussion

In this study, we have conducted a synthesis of the ecological information from the abstracts of 112,830 research articles spanning the published literature on biological pest control. The resulting database of pest herbivores and their natural enemies (DAPHNE) collates evidence for any ecological role or interaction to offer a more nuanced and holistic picture of a species’ ecology. As such, DAPHNE follows calls for evidence-based data syntheses in ecology and conservation (Sutherland et al., 2004), and provides a realistic, large-scale demonstration of how LLMs may be used to automate large parts of the evidence synthesis pipeline (Jaffer et al., 2025). Continually updating DAPHNE as new literature is published is technically trivial, and would pave the way towards ‘living’ syntheses (Berger-Tal et al., 2024), which may continue to evolve and improve as new information is added or previous data entries are corrected. Whilst it is certainly possible to have synthesised DAPHNE manually, the time expenditure demanded to analyse text at the scale conducted here would be prohibitively large in many contexts. Given sufficient computational resources, the approach outlined here could easily be scalable to analyse millions of scientific abstracts.

By extensively validating our dataset against a manually labelled training and test set, and by comparing results with other sources, this study has strived to follow best practices in the rapidly evolving domain of LLM-assisted text mining (Mammides and Papadopoulos, 2024; Moorthy et al., 2025; Scheepens et al., 2024), the results of which demonstrate that the information was extracted with high accuracy for the majority of data columns (Table 2). Indeed, comparing DAPHNE to other sources has revealed that the database is largely in agreement with previous classifications (Fig. 5), and is capable of yielding reliable evidence for interactions and classifications (e.g., pest status or pest regulation), some of which appear to have been overlooked in other sources. Moreover, we show that the reliability of the extractions in DAPHNE may be improved by filtering the database to exclude data entries with low evidence counts (Fig. 5).

At the same time, our results suggest that there is substantial variation in the reliability with which different pieces of information were extracted. For example, whilst the absence of biological control provision (the data column ‘Biocontrol’) was extracted with a precision of 99.0%, biocontrol as a true label was extracted with a much lower precision of 65.7%. This means that across the whole dataset, a false negative biocontrol label should be expected in about one out of 100 extractions, whilst a false positive should be expected in about one out of three cases. Despite relatively low precision, multiple independent sources evidencing biocontrol may nevertheless provide a strong indication for biocontrol provision, but it suggest that care is warranted when using DAPHNE to make conclusions based on low evidence-counts, particularly with regards to biocontrol provision (Fig. S12) and pest importance (Fig. S9). Whilst it is possible that our definition of biocontrol (evidenced if the natural enemy has been shown to exert control of the herbivore or is successfully used as a biological control agent of the herbivore) was not exact enough or too ambiguous, the quality and reliability of the extracted information is invariably limited by the clarity of the text, and ambiguous language, transcription mistakes and dubious translations can increase the likelihood of imprecise or erroneous extractions. Where the text is clearly written and unambiguous, however, extractions tend to be accurate (see Figs. S3, S4 and S5 for example extractions).

Whilst this study has demonstrated how LLMs can be utilised to automate a large part of the evidence synthesis pipeline, a significant effort is nevertheless demanded to ensure that the extracted information is both reliable and useful. This demands that considerable time is invested in manually labelling and validating a test set (as well as a ‘training’ set, if any form of prompt fine-tuning is adopted). For certain forms of data, direct comparisons of manual labels and model extractions may not be sensible (e.g., taxonomic terms, which are often misspelled and may be ambiguous), requiring further disambiguation (e.g., into minor and major mismatches, as done here). Furthermore, we highlight that a certain degree of post-processing of the LLM-generated output was required, including taxonomic corrections and harmonisation. These post-processing and manual validation steps can be time-consuming, but are important to ensure that a human stays in the loop and that risks of harm posed by LLM usage in research are minimised (Mammides and Papadopoulos, 2024; Weidinger et al., 2022).

Moreover, we acknowledge that DAPHNE exhibits considerable research bias towards the United States and China, given that 18.4% of publications (34,860) collected from Scopus originated from the United States and 9.6% (18,118) from China (Fig. 1). This bias is likely to be the result of analysing solely English-language studies, and could be ameliorated by filtering the dataset based on publication location. As most LLMs support a wide range of written languages, future studies may also extract data from literature in languages other than English (Berdejo-Espinola et al., 2025b; Liu et al., 2026), ensuring that non-English literature does not get lost in global ecological analyses (Amano et al., 2016, 2021; Konno et al., 2020), albeit with the caveat that model validation would have to account for multilingual literature.

We would like to caveat that despite our best efforts to validate the extracted data, only 0.5% of the interactions in DAPHNE have been manually verified and, where necessary, corrected. DAPHNE is also not an exhaustive synthesis and important interactions may be missing from the database and/or the literature that was analysed. Thus, data should be used carefully, and we advise that any interaction used should be manually checked and scrutinised, particularly when the interaction is evidenced by very few sources. Interactions evidenced by a large number of sources are expected to be more robust. Nevertheless, extractions should be understood as potential evidence for an interaction or a classification. In this way, we expect DAPHNE to be a helpful tool in gathering potential evidence for a wide range of use-cases, a few examples of which are detailed below.

### 4.1 Use cases

#### 4.1.1 Identification of important pests of host plants and their natural enemies

As detailed in Section 2.6, by summing evidence counts across the database, DAPHNE can be filtered to interactions with a minimum evidence count (e.g., nEID*≥* 5), or a selection of top *n* most-important crop pests (ranked on nEID, nPest or nMajor) and their top *m* most-important natural enemies (ranked on nEID, nImportant or nBiocontrol) to restrict the database to more reliable information (although evidence counts will to some extent reflect research bias). In this manner, DAPHNE can help identify potentially important pests of crops and other plants, their natural enemies and/or biocontrol agents.

#### 4.1.2 Identification of invasive species

DAPHNE contains 7112 interactions of between 759 unique herbivore taxa and their host plants where the herbivore was extracted to be invasive in a country or region. These data can aid in the identification of invasive (or potentially invasive) herbivores for invasive species research (e.g., Blackburn et al., 2014; Seebens et al., 2025).

#### 4.1.3 Identification of pathogen vectors

DAPHNE contains 4594 interactions of between 653 unique taxa (571 herbivores and 82 natural enemies) and their host plants or prey where the taxon was extracted to be a vector of a pathogen (e.g., a bacterium, virus, fungus, or otherwise pathogen-carrying). These data can aid in the identification of vectors (or potential vectors) of pathogens for host–pathogen and infectious disease research and data synthesis (Carlson et al., 2022; Stephens et al., 2017).

#### 4.1.4 Trophic network analysis

By extracting any available evidence for any interaction of interest, the data synthesised in DAPHNE can give rise to complex ecological networks (Fig. 3). Such networks can provide a basis for the analysis of ecological systems such as predator–prey or host–pathogen networks (Delmas et al., 2019; Guimaraes Jr, 2020; Runghen et al., 2021). The support of each interaction, as well as the LLM reasoning steps, can easily be interrogated by querying the database or even by visualising the entire network for a particular host-plant of interest (Fig. 6). To aid in the latter, DAPHNE can be used interactively for a selection of common crops and plantation tree species through a hosted Shiny web application (Scheepens, 2026a).

**Figure 6:**
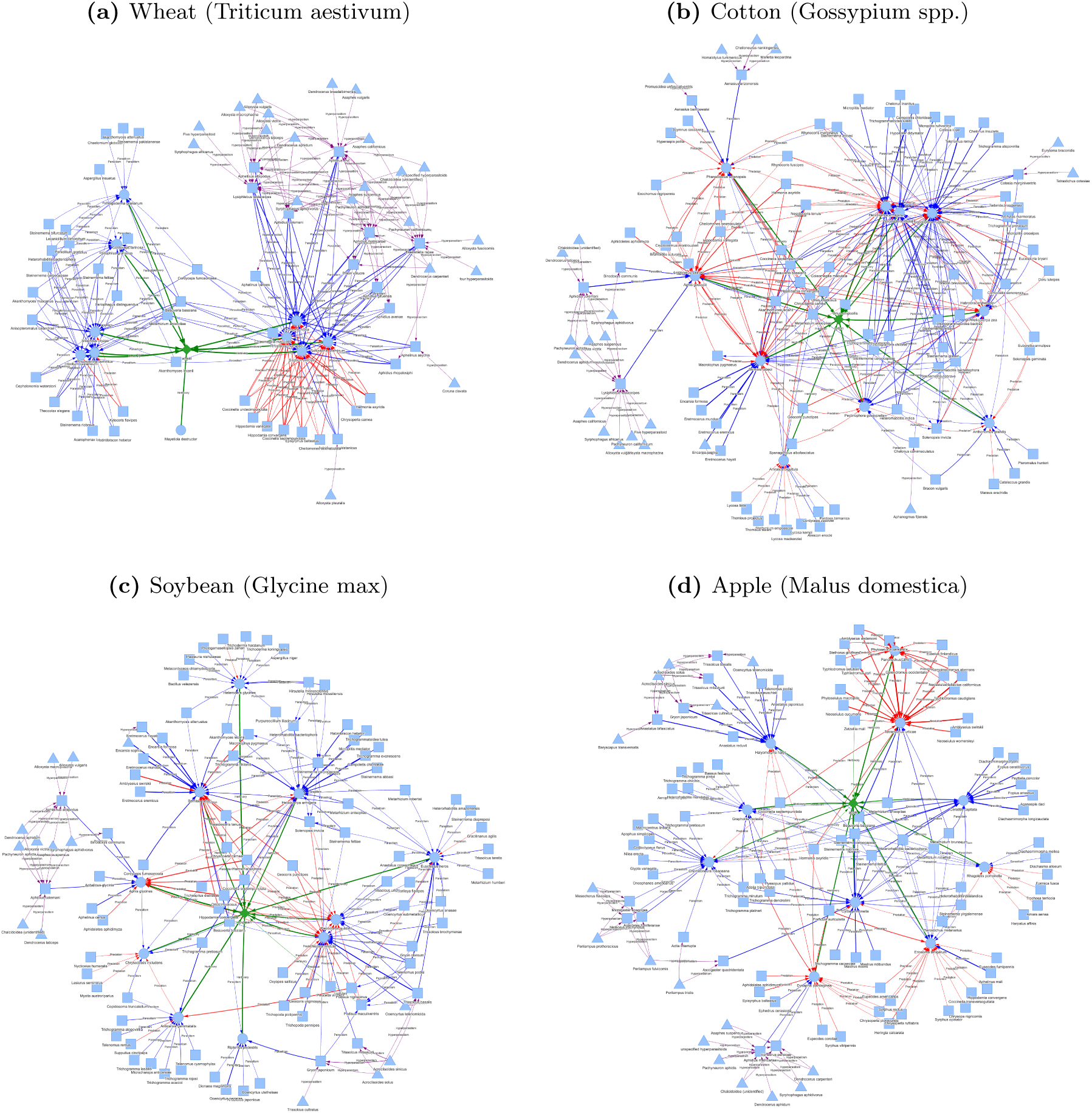
The interaction networks of ten globally important pests of (a) wheat, (b) cotton, (c) soybean and (d) apple (ranked on the number of abstracts evidencing the herbivore to be a pest of ‘major’ importance) and each pest’s ten most important natural enemies (ranked on the number of abstracts evidencing the species to be an ‘important’ natural enemy). Natural enemies (squares) interact with herbivorous hosts (circles) or other natural enemies through parasitism (blue arrows) or predation (red arrows). Hyperparasitoids (triangles) predating on parasitoids are plotted as purple arrows. Arrow widths are plotted (logarithmically) proportional to the number of abstracts evidencing the interaction. These networks can be viewed interactively through a hosted Shiny App (Scheepens, 2026a).

#### 4.1.5 Literature search and discovery

Beyond direct usage of the extracted data, DAPHNE maybe used as a search tool of the published pest control literature. For example, the database can be used to easily collect all published material in the database associated with a particular taxon, taxonomic group or interaction of interest, which can help researchers find potential candidate publications for further screening on a particular subject as it relates to pest control. The initial screening phase of the literature review process can be very time-consuming, which is where DAPHNE can offer significant support.

## 5 Conclusion

In this chapter, we developed DAPHNE, a large-scale, evidence-based synthesis of herbivorous pests, their host plants and their natural enemies from the published biological-control literature. By applying a pre-trained large language model to 112,830 scientific abstracts, we extracted 175,359 interactions involving 16,881 animal taxa resolved at the species level together with information on pest importance, biological control, invasiveness, pathogen vectoring and associated plants and industries. DAPHNE moves beyond assigning species to fixed functional groups by retaining evidence for individual ecological roles and interactions, allowing taxa to be represented differently across ecological contexts. As such, the database provides a structured, evidence-based resource for ecological interactions that are otherwise dispersed across a vast and largely unstructured literature. This framework is highly scalable and provides a major step towards continually evolving, ‘living’ syntheses of ecological information (Berger-Tal et al., 2024; Jaffer et al., 2025).

Validation showed that most information was extracted with high accuracy, with precision and recall generally exceeding 90%, and comparisons with other data sources showed a high level of agreement. These results demonstrate that carefully designed LLM workflows can substantially expand the scale of ecological evidence synthesis. At the same time, DAPHNE is not an exhaustive synthesis and important species or interactions may be missing from the database. Furthermore, only a small fraction of the complete database has been manually verified, which means that records should be manually checked and treated as candidate evidence. DAPHNE facilitates this by linking every piece of evidence back to the associated publication, along with the model’s own reasoning steps. Despite these limitations, DAPHNE provides a level of detail and magnitude far exceeding what could have been achieved with manual synthesis alone, and may help support the identification of important pests and natural-enemy relations, trophic network analyses, invasive species research, and literature search and discovery.

## 6 Data and code availability

All code and data used to create DAPHNE and produce the figures for this manuscript is available at https://github.com/dscheepens/DAPHNE-database-of-herbivores-and-natural-enemies, which is archived permanently in Zenodo (Scheepens, 2026b).

## Notes

https://github.com/dscheepens/DAPHNE-database-of-herbivores-and-natural-enemies

https://zenodo.org/records/24266976

### Competing Interest Statement

The authors have declared no competing interest.

### Summary of Updates

This version updates DAPHNE (v1.1, doi:10.5281/zenodo.24266976) after bug fixes to post-processing. Verbatim species names, taxonomic harmonisation with GBIF, and the matching of host plants to crops are corrected. Duplicate and placeholder records are removed. The database now contains 175,359 interactions and 16,881 species-level taxa. All figures and the EPPO comparison have been regenerated, and minor numerical errors in the text have been corrected. The main conclusions are unchanged.

### Funding

- Natural Environment Research Council: NE/S007229/1

