## Supplementary Information for "DAPHNE: A global database of pest herbivores and their natural enemies"

#### Contents

**A Equations** **2**

**B Tables** **3**

**C Figures** **13**

#### A Equations

$$\begin{aligned}
\text{PC} &= \frac{TP + TN}{TP + FP + TN + FN} \\
\text{Precision} &= \frac{TP}{TP + FP} \\
\text{Recall} &= \frac{TP}{TP + FN} \\
\text{F1-score} &= 2 \cdot \frac{\text{Precision} \cdot \text{Recall}}{\text{Precision} + \text{Recall}}
\end{aligned} \tag{1}$$

$$\begin{aligned}
TP &: \text{True Positives} \\
FP &: \text{False Positives} \\
TN &: \text{True Negatives} \\
FN &: \text{False Negatives}
\end{aligned}$$

### B Tables

**Table S1:** Complete descriptions of all data columns, as provided to the LLM in JSON format.

| Column | Data Type | JSON column description |
|---|---|---|
| Phylum | `string` | Taxonomic phylum of the species. |
| Class | `string` | Taxonomic class of the species. |
| Order | `string` | Taxonomic order of the species. |
| Family | `string` | Taxonomic family of the species. |
| Genus | `string` | Taxonomic genus of the species. |
| Species | `string` | Taxonomic species name (Latin binomial) or 'spp.' |
| Synonyms | List(`string`) | Any latin synonyms of this species stated in the text. These must be latin binomials, not common names. |
| Common Name | `string` | Only if the species has a common name associated with it in the text, state it here. Example: The common name of the sugarcane froghopper (Monecphora saccharina) is "The sugarcane froghopper". |
| (H) Feeding Mode | `categorical` [herbivory, frugivory, granivory, gall-formation] | Select if the species is a herbivore through granivory (seed predation), frugivory (fruit consumption), gall formation, or simply herbivory (default; any other form of herbivory, or unspecified herbivory). |
| (H) Host Plants | List(`string`) | (For each host plant) Plant or crop mentioned as host of the herbivorous species or as affected by it (add latin binomial if possible). This needs to be the plant specifically stated in the text to be affected or used as host, and should not be inferred from the herbivore's common name (e.g., if a herbivore called the "leek moth" is solely stated to be a pest of onions, then the host plant is onion, not leek). If a herbivorous species is stated to occur on a particular plant, or within a particular plantation/crop field, then it may be inferred to be a herbivore of this plant/crop. If the herbivore is stated or implicated as a general pest of agriculture, forestry or ornamentals without any particular host plants, then simply state 'agricultural crops', 'forestry' or 'ornamentals', respectively. Only select plants that are stated in the main abstract text - ignore plants that are only mentioned in the keywords. |
| (H) Is Pest | List(`boolean`) | (For each host plant) Is the herbivore a pest of this plant (True or False)? The species must either be mentioned explicitly as a pest of this plant, or at least as being the target of biological control (e.g., target of biocontrol study, pesticide application, etc.), or being the cause of destruction, infestation, commercial damage, etc., in relation to this plant. Pay close attention to the host crop or plant: If this is a weed or invasive, then the herbivore cannot be a pest of this plant; it can only be a pest if it affects plants that have commercial purposes in agriculture or forestry. |
| (H) Pest Importance | List(`categorical`) [minor, major] | (For each host plant) If the herbivore is a pest of this plant, is it of minor or major importance? Minor: The pest is stated to be of little economic importance or is only a nuisance and does not require biocontrol measures. Major: The herbivore is mentioned to be an important pest (e.g., highly destructive, commercially relevant, dominant, widespread pest, etc.), or is the intended target of biological control (e.g., target of a biocontrol or pesticide study) with regards to this plant/crop. Select 'NA' if nothing to this effect is mentioned or suggested in the text. |

Continued on next page

| Column | Data Type | JSON column description |
|---|---|---|
| (H) Is BCA | List(`boolean`) | (For each host plant) If the plant is a weed or is invasive, is the herbivore stated to be a biological control agent of the plant, or a promising/likely candidate for biological control (True or False)? |
| (H) Associated Industries | List(`categorical`) [arable crops, greenhouse production, pasture, fruit plantations, forestry and timber production, stored grains, ornamental and horticultural plants, unspecified agriculture, other] | List of any industries that are related to the host plants or crops of this herbivore, or are implied/mentioned in the text. If none of these industries are mentioned explicitly, choose whichever industries are most applicable to the host plants/crops of this herbivore. Choose 'unspecified agriculture' if none apply and agriculture is only mentioned in a general sense and no specific host plants are mentioned, and choose 'other' if it's an industry other than agriculture or forestry. Choose NA if no information about this is provided in the text whatsoever. |
| (H) Invasive In | List(`string`) | If this herbivore is explicitly described with the term 'invasive', in which countries or regions is it stated to be invasive? |
| (H) Pathogen Vector | List(`string`) | If the herbivore is stated to be a vector of a pathogen or disease, state the pathogen(s) and/or disease(s) here. |
| (NE) Feeding Mode | `categorical` [predation, parasitism] | Select if the species is a natural enemy through predation or parasitism (which includes parasitoidism). This may be inferred from the taxonomy (e.g. families of parasitoid wasps or predators). If not mentioned in the text or unclear, select NA. |
| (NE) Prey Name | List(`string`) | (For each prey) Name of the herbivorous species (binomial if possible) that is preyed upon or used as host by the natural enemy. These may also be groups of species (e.g. bark beetles, lepidopteran stemborers, etc., if only stated as such in the text). Look out for mentions of consumption: If a natural enemy is stated to consume a herbivorous species, this is evidence for the species being a natural enemy of the species. |
| (NE) Important Enemy | List(`boolean`) | (For each prey) Is the natural enemy stated to be an important natural enemy of this herbivorous prey/host (True or False)? Select True if the natural enemy is mentioned to be an important, dominant, frequent, common (etc.) natural enemy, is noted to exert important/effective/significant suppression/predation/parasitism, makes up a large proportion of the predation/parasitism of this herbivore, or has the potential to control/reduce/suppress the herbivore. Note: If the predation/parasitism of the natural enemy is found to be comparable to (or not found to be significantly different from) another species that is held to be an important natural enemy, then both species are important natural enemies. |
| (NE) Biocontrol | List(`boolean`) | (For each prey) Does the text explicitly state that the natural enemy provides biological control of this herbivorous prey/host (True or False)? Important: Select True ONLY if the natural enemy has been shown to provide/exert control of this herbivore or is successfully used as a biological control agent of this herbivore. If the text simply states that the natural enemy is a biological control agent of the herbivore, then assume that this means that it is successful - however, if the text mentions that the species was subsequently found not to provide effective control, then Biocontrol should be False. Important: If a natural enemy is mentioned to have 'potential' to control a herbivorous pest, then this makes it an important natural enemy of the herbivore (Important Enemy is True) - however, it does not suffice for Biocontrol to be True. Biocontrol should only be True if the natural enemy is found to effectively control the herbivorous pest, or if it is used or has been used successfully as a biological control agent. |
| (NE) Associated Plants | `string` | List of any plants associated with this natural enemy (e.g., where it is found to occur) or its herbivorous prey/hosts of (e.g., the affected crops, fruits, trees, flowers, etc.). |

Continued on next page

| Column | Data Type | JSON column description |
|---|---|---|
| (NE) Associated Industries | List(`categorical`) [arable crops, greenhouse production, pasture, fruit plantations, forestry and timber production, stored grains, ornamental and horticultural plants, unspecified agriculture, other] | List of any industries that are related to the host plants or crops associated with this natural enemy and/or its herbivorous prey/hosts. If none of these industries are mentioned explicitly, choose whichever industries are most applicable to the plants/crops associated with this natural enemy or its prey/hosts. Choose 'unspecified agriculture' if none apply and agriculture is only mentioned in a general sense and no specific host plants are mentioned, and choose 'other' if it's an industry other than agriculture or forestry. Choose NA if no information about this is provided in the text whatsoever. |
| (NE) Hyperparasitoids | List(`string`) | If the natural enemy is a parasitoid and the text mentions any hyperparasitoids parasitising this natural enemy, then list these hyperparasitoids here. |
| (NE) Invasive In | List(`string`) | If this natural enemy is explicitly described with the term 'invasive', in which countries or regions is it stated to be invasive? |
| (NE) Pathogen Vector | List(`string`) | If this natural enemy is a natural enemy of a herbivore due to vectoring a symbiotic bacterium/pathogen/disease that leads to mortality of the herbivorous prey/hosts, state it here. |

**Table S2:** Validation of model performance across all extracted data columns (variables). For taxonomy, the table shows proportion-correct (PC) scores, which were computed counting all mismatches as errors ($PC_{exact}$) and counting only major mismatches as errors ($PC_{approx}$). The latter was also computed separately for taxonomy that was directly available in the text ($PC^{extracted}_{approx}$) and taxonomy that was not available in the text and thus had to be 'inferred' ($PC^{inferred}_{approx}$). $N$ designates the total number of labels (also known as support). For the remaining columns, the table shows PC scores, precision (Pre), recall (Rec) and F1-scores.

| Column | $PC_{exact}$ | | $PC_{approx}$ | | $PC^{extracted}_{approx}$ | | $PC^{inferred}_{approx}$ | | $N$ | |
|---|---|---|---|---|---|---|---|---|---|---|
| | Train | Test | Train | Test | Train | Test | Train | Test | Train | Test |
| Phylum | 95.8 | 94.8 | 95.8 | 94.8 | 100 | 100 | 95.3 | 94.0 | 593 | 211 |
| Class | 99.7 | 95.7 | 99.7 | 95.7 | 100 | 100 | 99.4 | 93.4 | 593 | 211 |
| Order | 99.7 | 99.1 | 99.7 | 99.1 | 100 | 100 | 98.7 | 97.8 | 593 | 211 |
| Family | 96.8 | 96.2 | 98.0 | 97.2 | 99.8 | 99.2 | 88.9 | 94.1 | 593 | 211 |
| Genus | 99.0 | 100 | 99.7 | 100 | 99.7 | 100 | - | - | 593 | 211 |
| Species | 98.1 | 100 | 98.8 | 100 | 98.8 | 100 | - | - | 593 | 211 |

| Column | PC | | Pre | | Rec | | F1 | | $N$ | |
|---|---|---|---|---|---|---|---|---|---|---|
| | Train | Test | Train | Test | Train | Test | Train | Test | Train | Test |
| Synonyms | 100 | 97.2 | 100 | 0.0 | 100 | NA | 100 | NA | 593 | 211 |
| Common Name | 82.3 | 96.7 | 35.2 | 94.0 | 100 | 100 | 52.1 | 96.9 | 593 | 211 |
| Species Type | 99.8 | 100 | 99.8 | 100 | 99.8 | 100 | 99.8 | 100 | 593 | 211 |
| Feeding Mode | 99.3 | 100 | 99.7 | 100 | 99.3 | 100 | 99.4 | 100 | 593 | 211 |
| (H) Host Plants | 96.0 | 95.9 | 97.7 | 97.7 | 97.7 | 97.7 | 97.7 | 97.7 | 247 | 145 |
| (H) Is Pest | 93.9 | 91.7 | 94.1 | 91.9 | 93.9 | 91.7 | 94.0 | 91.8 | 247 | 145 |
| (H) Pest Importance | 92.7 | 93.1 | 93.1 | 93.3 | 92.7 | 93.1 | 92.8 | 93.1 | 247 | 145 |
| (H) Is BCA | 95.1 | 96.6 | 95.1 | 96.7 | 95.1 | 96.6 | 95.1 | 96.6 | 247 | 145 |
| (NE) Herbivore Prey | 98.9 | 91.9 | 99.2 | 96.1 | 99.6 | 94.3 | 99.4 | 95.2 | 533 | 124 |
| (NE) Important Enemy | 93.6 | 87.9 | 93.7 | 89.2 | 93.6 | 87.9 | 93.6 | 88.4 | 533 | 124 |
| (NE) Biocontrol | 96.8 | 83.9 | 97.4 | 87.6 | 96.8 | 83.9 | 97.0 | 84.5 | 533 | 124 |
| (NE) Hyperparasitoids | 100 | 100 | 100 | 100 | 100 | 100 | 100 | 100 | 447 | 101 |
| (NE) Associated Plants | 99.1 | 96.2 | 98.9 | 95.1 | 100 | 100 | 99.5 | 97.5 | 461 | 106 |
| Invasive In | 100 | 99.1 | 100 | 75.0 | 100 | 100 | 100 | 85.7 | 598 | 213 |
| Vectors | 99.7 | 100 | 100 | 100 | 80.0 | 100 | 88.9 | 100 | 593 | 211 |
| Associated Industries | 86.0 | 97.0 | 89.8 | 97.0 | 86.0 | 96.9 | 87.2 | 96.8 | 657 | 234 |

**Table S3:** Patterns used to string-match crop names with crops and plants in the DAPHNE database.

<table>
<thead>
<tr><th>Crop Name</th><th>Pattern</th><th>Group</th><th>Class</th></tr>
</thead>
<tbody>
<tr><td>wheat</td><td>\b(wheats?|triticum)\b</td><td>Cereals</td><td>Wheat</td></tr>
<tr><td>maize</td><td>\b(maize|corns?|zea mays)\b</td><td>Cereals</td><td>Maize</td></tr>
<tr><td>rice</td><td>\b(rice|oryza)\b</td><td>Cereals</td><td>Rice</td></tr>
<tr><td>barley</td><td>\b(barleys?|hordeum)\b</td><td>Cereals</td><td>Barley</td></tr>
<tr><td>sorghum</td><td>\b(sorghums?|great millet|indian millet)\b</td><td>Cereals</td><td>Sorghum</td></tr>
<tr><td>millet</td><td>\b((?<!pearl )(?<!great )(?<!indian )millet|panicum|setaria|eleusine|pennisetum|digitaria)\b</td><td>Cereals</td><td>Millets</td></tr>
<tr><td>rye</td><td>\b(rye|secale)\b</td><td>Cereals</td><td>Rye</td></tr>
<tr><td>oats</td><td>\b(oats?|avena)\b</td><td>Cereals</td><td>Oats</td></tr>
<tr><td>mixed cereals</td><td>\b(cereals?|grains?)\b</td><td>Cereals</td><td>Other cereals</td></tr>
<tr><td>cabbage</td><td>\b(cabbages?|brassica oleracea)\b</td><td>Vegetables</td><td>Leafy vegetables</td></tr>
<tr><td>lettuce</td><td>\b(lettuces?|lactuca sativa)\b</td><td>Vegetables</td><td>Leafy vegetables</td></tr>
<tr><td>spinach</td><td>\b(spinach|spinacia)\b</td><td>Vegetables</td><td>Leafy vegetables</td></tr>
<tr><td>tomato</td><td>\b(tomatoes|tomato|solanum lycopersicum)\b</td><td>Vegetables</td><td>Fruit-bearing vegetables</td></tr>
<tr><td>eggplant</td><td>\b(eggplants?|aubergines?|solanum melongena)\b</td><td>Vegetables</td><td>Fruit-bearing vegetables</td></tr>
<tr><td>cucumber</td><td>\b(cucumbers?|cucumis sativus)\b</td><td>Vegetables</td><td>Fruit-bearing vegetables</td></tr>
<tr><td>melon</td><td>\b(melons?|cucumis melo|cantaloupes?)\b</td><td>Vegetables</td><td>Fruit-bearing vegetables</td></tr>
<tr><td>pumpkin</td><td>\b(pumpkins?|squash|gourds?|cucurbita)\b</td><td>Vegetables</td><td>Fruit-bearing vegetables</td></tr>
<tr><td>carrot</td><td>\b(carrots?|daucus carota)\b</td><td>Vegetables</td><td>Root vegetables</td></tr>
<tr><td>onion</td><td>\b(onions?|allium cepa)\b</td><td>Vegetables</td><td>Bulb vegetables</td></tr>
<tr><td>garlic</td><td>\b(garlic|allium sativum)\b</td><td>Vegetables</td><td>Bulb vegetables</td></tr>
<tr><td>banana</td><td>\b(bananas?|musa)\b</td><td>Fruit</td><td>Tropical fruit</td></tr>
<tr><td>plantain</td><td>\b(plantains?|musa paradisiaca)\b</td><td>Fruit</td><td>Tropical fruit</td></tr>
<tr><td>mango</td><td>\b(mangos?|mangoes?|mangifera indica)\b</td><td>Fruit</td><td>Tropical fruit</td></tr>
<tr><td>papaya</td><td>\b(papayas?|carica papaya)\b</td><td>Fruit</td><td>Tropical fruit</td></tr>
<tr><td>pineapple</td><td>\b(pineapples?|ananas)\b</td><td>Fruit</td><td>Tropical fruit</td></tr>
<tr><td>avocado</td><td>\b(avocados?|persea americana)\b</td><td>Fruit</td><td>Tropical fruit</td></tr>
<tr><td>orange</td><td>\b(oranges?|citrus sinensis)\b</td><td>Fruit</td><td>Citrus</td></tr>
<tr><td>lemon</td><td>\b(lemons?|citrus limon)\b</td><td>Fruit</td><td>Citrus</td></tr>
<tr><td>lime</td><td>\b(limes?|citrus aurantiifolia)\b</td><td>Fruit</td><td>Citrus</td></tr>
<tr><td>grapefruit</td><td>\b(grapefruits?|citrus paradisi)\b</td><td>Fruit</td><td>Citrus</td></tr>
<tr><td>apple</td><td>\b(apples?|malus domestica)\b</td><td>Fruit</td><td>Pome fruit</td></tr>
<tr><td>pear</td><td>\b(pears?|pyrus)\b</td><td>Fruit</td><td>Pome fruit</td></tr>
<tr><td>peach</td><td>\b(peaches?|peach|prunus persica)\b</td><td>Fruit</td><td>Stone fruit</td></tr>
<tr><td>plum</td><td>\b(plums?|prunus domestica)\b</td><td>Fruit</td><td>Stone fruit</td></tr>
<tr><td>cherry</td><td>\b(cherries|cherry|prunus avium)\b</td><td>Fruit</td><td>Stone fruit</td></tr>
<tr><td>grape</td><td>\b(grape(?!fruit)s?|vitis)\b</td><td>Fruit</td><td>Grapes</td></tr>
</tbody>
</table>

Continued on next page

**Table S3 – Continued from previous page**

<table>
<tr><th>Crop Name</th><th>Pattern</th><th>Group</th><th>Class</th></tr>
<tr><td>almond</td><td>\b(almonds?|prunus dulcis)\b</td><td>Fruit</td><td>Nuts</td></tr>
<tr><td>cashew</td><td>\b(cashews?|anacardium occidentale)\b</td><td>Fruit</td><td>Nuts</td></tr>
<tr><td>walnut</td><td>\b(walnuts?|juglans)\b</td><td>Fruit</td><td>Nuts</td></tr>
<tr><td>soybean</td><td>\b(soybeans?|soy\s?beans?|glycine max)\b</td><td>Oilseed crops</td><td>Soya beans</td></tr>
<tr><td>groundnut</td><td>\b(groundnuts?|peanuts?|arachis hypogaea)\b</td><td>Oilseed crops</td><td>Groundnuts</td></tr>
<tr><td>rapeseed</td><td>\b(rapeseeds?|canola|brassica napus)\b</td><td>Oilseed crops</td><td>Rapeseed</td></tr>
<tr><td>sunflower</td><td>\b(sunflowers?|helianthus annuus)\b</td><td>Oilseed crops</td><td>Sunflower</td></tr>
<tr><td>sesame</td><td>\b(sesame|sesamum indicum)\b</td><td>Oilseed crops</td><td>Sesame</td></tr>
<tr><td>linseed</td><td>\b(linseed|flax|linum usitatissimum)\b</td><td>Oilseed crops</td><td>Linseed</td></tr>
<tr><td>mustard</td><td>\b(mustard|brassica)\b</td><td>Oilseed crops</td><td>Mustard</td></tr>
<tr><td>coconut</td><td>\b(coconuts?|cocos nucifera)\b</td><td>Oilseed crops</td><td>Permanent oil crops</td></tr>
<tr><td>oil palm</td><td>\b(oil\s?palms?|elaeis guineensis)\b</td><td>Oilseed crops</td><td>Permanent oil crops</td></tr>
<tr><td>olive</td><td>\b(olives?|olea europaea)\b</td><td>Oilseed crops</td><td>Permanent oil crops</td></tr>
<tr><td>potato</td><td>\b((?<!sweet )potatoes|(?<!sweet )potato|solanum tuberosum)\b</td><td>Roots & tubers</td><td>Potatoes</td></tr>
<tr><td>sweet potato</td><td>\b(sweet potatoes|sweet potato|ipomoea batatas)\b</td><td>Roots & tubers</td><td>Sweet potatoes</td></tr>
<tr><td>cassava</td><td>\b(cassava|manihot esculenta|tapioca)\b</td><td>Roots & tubers</td><td>Cassava</td></tr>
<tr><td>yam</td><td>\b(yams?|dioscorea)\b</td><td>Roots & tubers</td><td>Yams</td></tr>
<tr><td>taro</td><td>\b(taro|colocasia esculenta)\b</td><td>Roots & tubers</td><td>Other roots</td></tr>
<tr><td>coffee</td><td>\b(coffees?|coffea arabica|coffea canephora)\b</td><td>Beverage crops</td><td>Coffee</td></tr>
<tr><td>tea</td><td>\b(teas?|camellia sinensis)\b</td><td>Beverage crops</td><td>Tea</td></tr>
<tr><td>cocoa</td><td>\b(cocoa|cacao|theobroma cacao)\b</td><td>Beverage crops</td><td>Cocoa</td></tr>
<tr><td>chili</td><td>\b(chili|chilies|capsicum|(?<!black )pepper(?!\s+spice))\b</td><td>Spice crops</td><td>Temporary spices</td></tr>
<tr><td>pepper spice</td><td>\b(black pepper|piper nigrum)\b</td><td>Spice crops</td><td>Permanent spices</td></tr>
<tr><td>ginger</td><td>\b(ginger|zingiber officinale)\b</td><td>Spice crops</td><td>Permanent spices</td></tr>
<tr><td>cinnamon</td><td>\b(cinnamon|cinnamomum)\b</td><td>Spice crops</td><td>Permanent spices</td></tr>
<tr><td>beans</td><td>(?<!soy)(?<!coffee\s)\bbeans?\b|\bphaseolus\b|\bvigna\b(?!\s+unguiculata)</td><td>Legumes</td><td>Beans</td></tr>
<tr><td>chickpea</td><td>\b(chickpeas?|cicer arietinum)\b</td><td>Legumes</td><td>Chickpeas</td></tr>
<tr><td>cowpea</td><td>\b(cowpeas?|vigna unguiculata)\b</td><td>Legumes</td><td>Cowpeas</td></tr>
<tr><td>lentil</td><td>\b(lentils?|lens culinaris)\b</td><td>Legumes</td><td>Lentils</td></tr>
<tr><td>pea</td><td>\b((?<!pigeon\s)peas?|pisum sativum)\b</td><td>Legumes</td><td>Peas</td></tr>
<tr><td>pigeon pea</td><td>\b(pigeon peas?|cajanus cajan)\b</td><td>Legumes</td><td>Pigeon peas</td></tr>
<tr><td>sugarcane</td><td>\b(sugarcane|sugar\s?cane|saccharum officinarum)\b</td><td>Sugar crops</td><td>Sugar cane</td></tr>
<tr><td>sugarbeet</td><td>\b(sugarbeet|sugar\s?beet|beta vulgaris)\b</td><td>Sugar crops</td><td>Sugar beet</td></tr>
<tr><td>sweet sorghum</td><td>\b(sweet sorghum)\b</td><td>Sugar crops</td><td>Other sugar crops</td></tr>
<tr><td>cotton</td><td>\b(cotton|gossypium)\b</td><td>Fibre crops</td><td>Cotton</td></tr>
<tr><td>jute</td><td>\b(jute|corchorus)\b</td><td>Fibre crops</td><td>Fibre crops</td></tr>
<tr><td>flax</td><td>\b(flax|linum)\b</td><td>Fibre crops</td><td>Fibre crops</td></tr>
</table>

Continued on next page

**Table S3 – Continued from previous page**

| Crop Name | Pattern | Group | Class |
|---|---|---|---|
| rubber | `\b(rubber|hevea brasiliensis)\b` | Industrial crops | Rubber |
| tobacco | `\b(tobacco|nicotiana)\b` | Industrial crops | Tobacco |
| alfalfa | `\b(alfalfa|medicago sativa)\b` | Fodder crops | Fodder |
| grass | `\b(grass|fodder|pasture)\b` | Fodder crops | Fodder |
| pine | `\b(pines?|pinus)\b` | Forest plantation | Softwood forest |
| douglas fir | `\b(douglas-fir|douglas fir|pseudotsuga menziesii)\b` | Forest plantation | Softwood forest |
| spruce | `\b(spruces?|picea)\b` | Forest plantation | Softwood forest |
| teak | `\b(teak|tectona grandis)\b` | Forest plantation | Hardwood forest |
| eucalyptus | `\b(eucalyptus|eucalypts?)\b` | Forest plantation | Hardwood forest |

**Table S4:** Comparison of DAPHNE with the global EPPO (European and Mediterranean Plant Protection Organization) database (EPPO, 2026) for a selection of crops and plantation trees. The table shows the number of hosts ($n$ Hosts) and major hosts ($n$ Major Hosts) as listed in EPPO. For each crop, the respective hosts and major hosts were matched with species extracted in DAPHNE as herbivores of this crop. Based on the LLM-extracted evidence, we then divided these species into pests (nPest>0) and major pests (nMajor>0). To determine how well EPPO hosts were captured in DAPHNE, we computed recall scores 1) between pests (nPest>0) and species classified as hosts or major hosts, 2) between major pests (nMajor>0) and major hosts, and 3) between pests (nPest>0) and major hosts (annotated with an asterisk in the table). The table shows that there is substantial variation between crops, but overall, species classified as hosts or major hosts in EPPO were captured as pests (nPest>0) with 93.1% recall, species classified as major hosts in EPPO were captured as major pests (nMajor>0) with 90.1% recall, and were captured as pests (nPest>0) with 97.7% recall. The table also shows the mean number of counts evidencing pest status (nPest) for both hosts and major hosts, as well as the mean number of EIDs (nEID) evidencing herbivory of any kind (i.e., irrespective of pest status). Overall, major hosts obtained marginally higher nEID and nPest counts than hosts, although there is a lot of variation between crops.

| Crop name | $n$ Hosts | $n$ Major Hosts | Recall (%) | | | Mean nPest | | Mean nEID | |
|---|---|---|---|---|---|---|---|---|---|
| | | | Hosts & Major Hosts | Major Hosts | Major Hosts* | Hosts | Major Hosts | Hosts | Major Hosts |
| wheat | 14 | 3 | 92.9 | 66.7 | 66.7 | 21.6 | 2.7 | 22.0 | 4.3 |
| maize | 52 | 26 | 96.2 | 92.3 | 100.0 | 13.2 | 110.5 | 15.0 | 113.3 |
| rice | 46 | 27 | 97.8 | 92.6 | 100.0 | 13.7 | 74.7 | 14.0 | 76.2 |
| barley | 10 | 3 | 70.0 | 100.0 | 100.0 | 12.3 | 5.3 | 13.9 | 5.3 |
| sorghum | 6 | 1 | 66.7 | 0.0 | 100.0 | 13.0 | 1.0 | 14.0 | 1.0 |
| millet | 9 | 0 | 66.7 | NaN | NaN | 4.9 | NaN | 9.0 | NaN |
| rye | 7 | 0 | 71.4 | NaN | NaN | 1.3 | NaN | 1.7 | NaN |
| oats | 6 | 0 | 66.7 | NaN | NaN | 2.0 | NaN | 2.7 | NaN |
| cabbage | 29 | 6 | 96.6 | 100.0 | 100.0 | 13.1 | 26.2 | 15.1 | 29.3 |
| lettuce | 9 | 2 | 88.9 | 50.0 | 100.0 | 2.7 | 1.5 | 3.3 | 2.5 |
| spinach | 2 | 0 | 100.0 | NaN | NaN | 1.0 | NaN | 2.0 | NaN |
| tomato | 67 | 26 | 91.0 | 92.3 | 100.0 | 18.5 | 57.0 | 19.9 | 58.3 |
| eggplant | 35 | 11 | 97.1 | 90.9 | 100.0 | 7.9 | 15.1 | 9.2 | 15.6 |
| cucumber | 26 | 5 | 96.2 | 80.0 | 80.0 | 9.9 | 18.8 | 11.2 | 21.8 |
| melon | 22 | 6 | 100.0 | 100.0 | 100.0 | 6.7 | 6.7 | 7.0 | 7.2 |
| pumpkin | 27 | 6 | 96.3 | 83.3 | 100.0 | 6.9 | 11.2 | 8.1 | 11.8 |
| carrot | 4 | 1 | 100.0 | 100.0 | 100.0 | 1.0 | 30.0 | 1.0 | 31.0 |
| onion | 20 | 4 | 100.0 | 100.0 | 100.0 | 4.6 | 12.8 | 5.1 | 13.0 |
| garlic | 7 | 1 | 100.0 | 100.0 | 100.0 | 3.5 | 5.0 | 3.7 | 5.0 |
| banana | 27 | 7 | 96.3 | 100.0 | 100.0 | 10.8 | 20.1 | 11.2 | 20.1 |

Continued on next page

**Table S4 – Continued from previous page**

| Crop name | $n$ Hosts | $n$ Major Hosts | Recall (%) | | | Mean nPest | | Mean nEID | |
|---|---|---|---|---|---|---|---|---|---|
| | | | Hosts & Major Hosts | Major Hosts | Major Hosts* | Hosts | Major Hosts | Hosts | Major Hosts |
| plantain | 10 | 3 | 90.0 | 100.0 | 100.0 | 3.1 | 3.3 | 3.3 | 3.3 |
| mango | 47 | 11 | 93.6 | 100.0 | 100.0 | 5.5 | 14.7 | 6.2 | 15.6 |
| papaya | 21 | 3 | 81.0 | 100.0 | 100.0 | 2.9 | 2.0 | 3.4 | 3.0 |
| pineapple | 3 | 1 | 100.0 | 0.0 | 100.0 | 6.0 | 1.0 | 6.0 | 1.0 |
| avocado | 34 | 11 | 94.1 | 81.8 | 100.0 | 4.1 | 8.3 | 4.3 | 8.6 |
| orange | 56 | 19 | 96.4 | 100.0 | 100.0 | 7.2 | 5.7 | 7.9 | 6.1 |
| lemon | 31 | 11 | 87.1 | 81.8 | 100.0 | 3.0 | 4.0 | 4.0 | 4.4 |
| lime | 13 | 1 | 100.0 | 100.0 | 100.0 | 2.7 | 1.0 | 2.8 | 1.0 |
| grapefruit | 27 | 8 | 92.6 | 87.5 | 100.0 | 3.7 | 4.9 | 3.9 | 5.0 |
| apple | 56 | 19 | 94.6 | 68.4 | 89.5 | 7.9 | 22.6 | 8.4 | 23.5 |
| pear | 35 | 4 | 97.1 | 100.0 | 100.0 | 6.4 | 19.2 | 6.7 | 19.2 |
| peach | 45 | 15 | 93.3 | 86.7 | 93.3 | 8.5 | 12.3 | 9.1 | 13.0 |
| plum | 24 | 5 | 91.7 | 100.0 | 100.0 | 2.6 | 5.4 | 3.1 | 6.2 |
| cherry | 29 | 10 | 89.7 | 90.0 | 90.0 | 2.5 | 18.2 | 2.8 | 18.8 |
| grape | 4 | 0 | 100.0 | NaN | NaN | 10.2 | NaN | 12.0 | NaN |
| almond | 17 | 3 | 94.1 | 66.7 | 100.0 | 2.4 | 4.3 | 2.5 | 4.3 |
| cashew | 15 | 3 | 93.3 | 100.0 | 100.0 | 1.9 | 6.0 | 2.0 | 6.0 |
| walnut | 15 | 2 | 86.7 | 100.0 | 100.0 | 7.1 | 5.0 | 8.0 | 5.5 |
| soybean | 27 | 9 | 96.3 | 100.0 | 100.0 | 14.6 | 76.2 | 17.3 | 79.4 |
| groundnut | 16 | 2 | 100.0 | 100.0 | 100.0 | 8.9 | 4.5 | 10.4 | 4.5 |
| rapeseed | 18 | 1 | 94.4 | 100.0 | 100.0 | 5.4 | 1.0 | 5.6 | 1.0 |
| sunflower | 10 | 1 | 70.0 | 100.0 | 100.0 | 3.6 | 1.0 | 5.4 | 1.0 |
| sesame | 5 | 0 | 80.0 | NaN | NaN | 2.0 | NaN | 2.6 | NaN |
| linseed | 2 | 0 | 100.0 | NaN | NaN | 1.0 | NaN | 2.5 | NaN |
| mustard | 28 | 1 | 96.4 | 100.0 | 100.0 | 16.7 | 27.0 | 19.2 | 30.0 |
| coconut | 24 | 16 | 95.8 | 87.5 | 100.0 | 1.2 | 20.3 | 1.5 | 20.6 |
| oil palm | 13 | 6 | 84.6 | 83.3 | 100.0 | 2.9 | 3.5 | 3.1 | 3.5 |
| olive | 12 | 1 | 100.0 | 100.0 | 100.0 | 8.5 | 295.0 | 8.7 | 298.0 |
| potato | 58 | 29 | 96.6 | 96.6 | 100.0 | 5.7 | 40.3 | 6.3 | 41.2 |
| sweet potato | 19 | 6 | 84.2 | 100.0 | 100.0 | 4.9 | 28.5 | 6.2 | 30.2 |
| cassava | 11 | 4 | 100.0 | 75.0 | 100.0 | 4.1 | 55.5 | 4.4 | 56.2 |
| yam | 1 | 0 | 100.0 | NaN | NaN | 4.0 | NaN | 5.0 | NaN |
| taro | 4 | 2 | 50.0 | 50.0 | 50.0 | 3.5 | 0.5 | 5.5 | 1.0 |
| coffee | 25 | 10 | 100.0 | 100.0 | 100.0 | 25.9 | 14.6 | 26.4 | 15.1 |

Continued on next page

**Table S4 – Continued from previous page**

| Crop name | $n$ Hosts | $n$ Major Hosts | Recall (%) | | | Mean nPest | | Mean nEID | |
|---|---|---|---|---|---|---|---|---|---|
| | | | Hosts & Major Hosts | Major Hosts | Major Hosts* | Hosts | Major Hosts | Hosts | Major Hosts |
| tea | 13 | 5 | 100.0 | 100.0 | 100.0 | 3.5 | 22.2 | 3.5 | 22.4 |
| cocoa | 25 | 16 | 92.0 | 81.2 | 93.8 | 1.8 | 6.9 | 2.0 | 7.1 |
| chili | 5 | 0 | 80.0 | NaN | NaN | 33.4 | NaN | 34.6 | NaN |
| pepper spice | 1 | 0 | 100.0 | NaN | NaN | 2.0 | NaN | 2.0 | NaN |
| ginger | 3 | 0 | 100.0 | NaN | NaN | 2.0 | NaN | 2.3 | NaN |
| beans | 6 | 0 | 83.3 | NaN | NaN | 25.0 | NaN | 27.3 | NaN |
| chickpea | 6 | 1 | 100.0 | 100.0 | 100.0 | 37.6 | 3.0 | 39.2 | 3.0 |
| cowpea | 18 | 1 | 88.9 | 100.0 | 100.0 | 4.8 | 117.0 | 6.2 | 117.0 |
| lentil | 1 | 0 | 100.0 | NaN | NaN | 1.0 | NaN | 1.0 | NaN |
| pea | 15 | 1 | 80.0 | 0.0 | 100.0 | 4.2 | 1.0 | 4.9 | 1.0 |
| pigeon pea | 17 | 6 | 76.5 | 100.0 | 100.0 | 15.3 | 8.3 | 16.9 | 8.5 |
| sugarcane | 35 | 17 | 100.0 | 88.2 | 100.0 | 15.7 | 19.1 | 15.9 | 19.6 |
| sugarbeet | 17 | 8 | 94.1 | 100.0 | 100.0 | 2.8 | 11.4 | 3.6 | 12.0 |
| cotton | 18 | 1 | 100.0 | 100.0 | 100.0 | 51.5 | 90.0 | 53.6 | 92.0 |
| flax | 2 | 0 | 100.0 | NaN | NaN | 1.0 | NaN | 2.5 | NaN |
| rubber | 9 | 1 | 100.0 | 100.0 | 100.0 | 1.4 | 3.0 | 1.4 | 3.0 |
| tobacco | 19 | 7 | 84.2 | 100.0 | 100.0 | 7.1 | 16.3 | 11.2 | 20.3 |
| alfalfa | 19 | 5 | 78.9 | 100.0 | 100.0 | 7.7 | 7.2 | 8.7 | 9.0 |
| pine | 26 | 2 | 100.0 | 100.0 | 100.0 | 29.6 | 17.5 | 30.5 | 24.0 |
| douglas fir | 11 | 6 | 100.0 | 100.0 | 100.0 | 1.4 | 8.5 | 1.4 | 9.2 |
| spruce | 12 | 3 | 83.3 | 0.0 | 33.3 | 19.0 | 0.3 | 19.3 | 1.3 |
| teak | 2 | 0 | 100.0 | NaN | NaN | 2.0 | NaN | 2.0 | NaN |
| eucalyptus | 11 | 4 | 100.0 | 100.0 | 100.0 | 23.4 | 8.2 | 25.0 | 9.2 |
| **overall** | **1471** | **436** | **93.1** | **90.1** | **97.7** | **59.4** | **62.0** | **64.3** | **66.7** |

#### C Figures

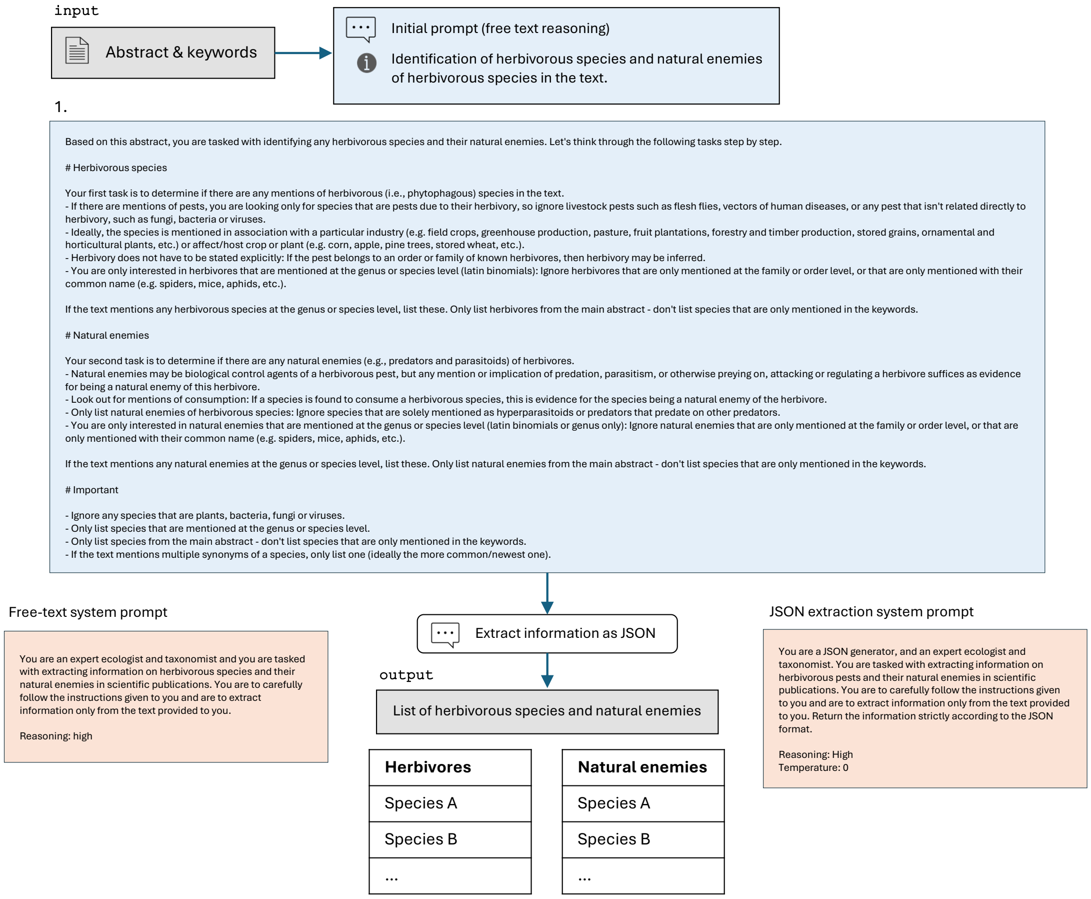


**Figure S1:** Prompt diagram, part 1: Extraction of lists of herbivores and their natural enemies from the abstract & keywords. The LLM (gpt-oss:120b) was first prompted to return a free-text response to a detailed, zero-shot prompt using the free-text system prompt. The free-text response was then provided as another prompt to a new instantiation of the LLM using the JSON extraction system prompt, prompting the model to return the text according to the specified JSON scheme.

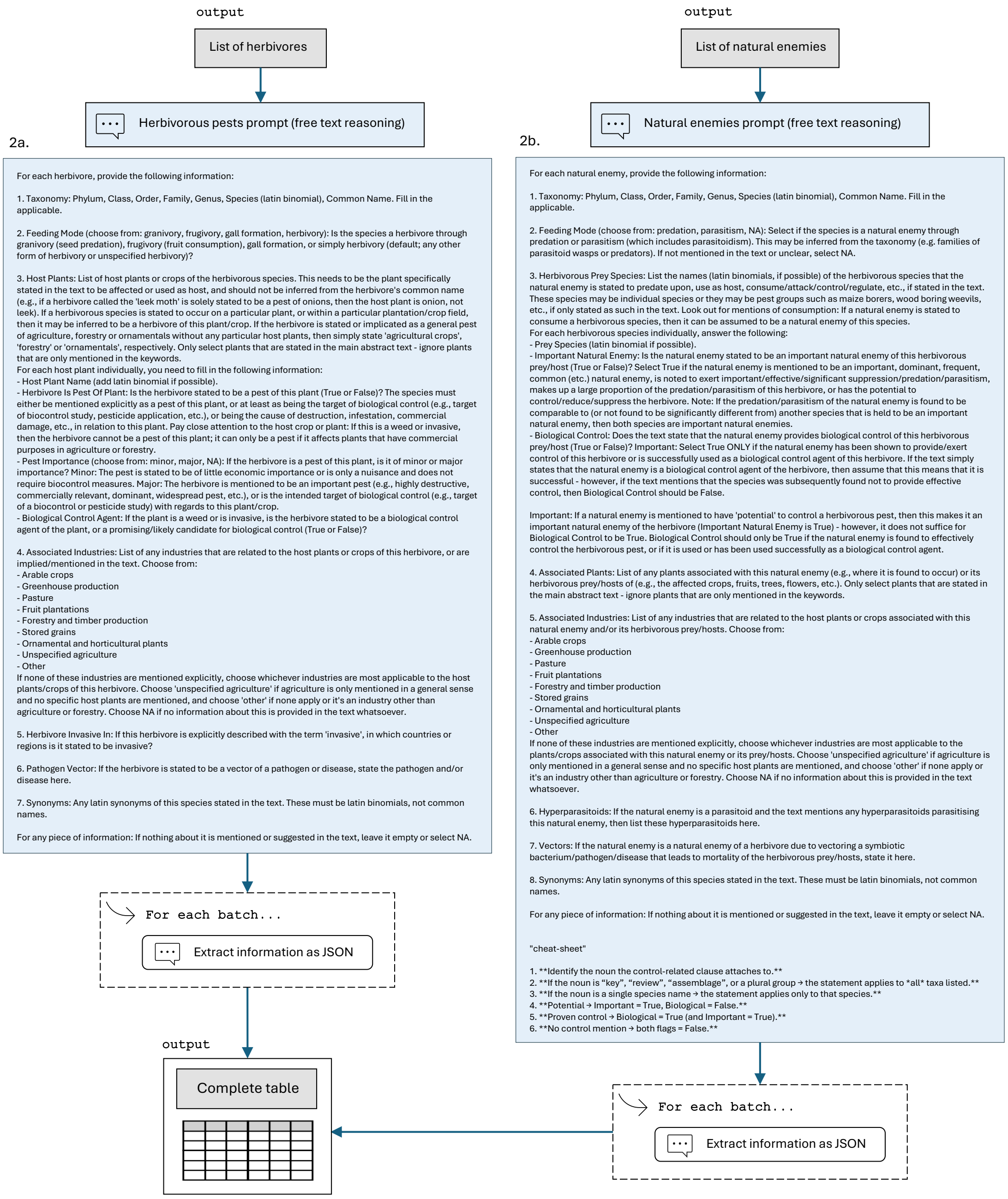


**Figure S2:** Prompt diagram, part 2: Further extraction of species-specific information. Using the extracted lists of herbivores and their natural enemies (Fig. S1), the LLM was prompted to return a free-text response to the respective prompt for one batch (five species) at a time. The LLM was then prompted again to return the free-text response according to the specified JSON scheme, and all JSON output was finally collected in a complete table. We found that allowing the LLM to first generate a free-text response improved its ability to reason about complicated and nuanced cases of pest status or biological control provision.

Life cycle and food consumption potential of the invasive terrestrial slug Meghimatium pictum (Stoliczka, 1873); Ciclo de vida e potencial de consumo alimentar da lesma terres
tre invasora Meghimatium pictum (Stoliczka, 1873). Abstract: The invasive chinese slug Meghimatium pictum (Stoliczka, 1873) (Stylommathophora: Philomycidae) is originally fr
om Asia, and it has been introduced in Latin American countries like Argentina and Brazil, where it is considered a critical horticultural pest. This species also became an interm
ediate host for the nematode Angiostrongylus costaricensis Morera & Cespedes, 1971 (Strongylida: Metastrongylidae), which can cause abdominal angiostrongyliasis in humans
when ingested molluscs or their mucus containing larvae released on fruit and vegetables. This research aimed to investigate the biological parameters of the life cycle of M. pic
tum and evaluate its food preference to understand the species' behavior and provide information on the choice of safer pest management and control methods. We observed t
hat 68 and 75% of the grouped and isolated slugs, respectively, survived 26 weeks (180 days) under laboratory conditions. In addition, the individuals kept isolated had higher bo
dy mass (2.8 ± 0.6 g), length (3.3 ± 0.8 cm), and width (0.37 ± 0.3 cm) than grouped specimens. We also found that M. pictum has indeterminate growth and an annual reproducti
ve cycle. Concerning food preference, slugs better accepted lettuce at different developmental stages (neonate, juvenile, and adult). Our study presents the first description of t
he M. pictum life cycle. We concluded that M. pictum has undefined biological parameters, which hampers its laboratory rearing. However, we also demonstrate its potential as
a pest for different horticultural crops, which will require the development of management strategies. © 2024 Elsevier B.V., All rights reserved.. Author keywords: Growth; Philomy
cidae; Reproduction; Terrestrial Gastropod.

| Subject | Object | Type | Feeding Mode | Is Pest | Pest Importance | Herbivore Is BCA | Associated Industries | Invasive In | Vectors | Important Enemy | Biocontrol | Associated Plants |
|---|---|---|---|---|---|---|---|---|---|---|---|---|
| Meghimatium pictum | Lactuca sativa | herbivore | herbivory | FALSE | NA | FALSE | ornamental and horticultural plants | Argentina, Brazil | Angiostrongylus costaricensis (abdominal angi ostrongyliasis) | NA | NA | NA |
| Meghimatium pictum | horticultural crops | herbivore | herbivory | TRUE | major | FALSE | ornamental and horticultural plants | Argentina, Brazil | Angiostrongylus costaricensis (abdominal angi ostrongyliasis) | NA | NA | NA |
| Angiostrongylus costaricensis | Meghimatium pictum | natural enemy | parasitism | NA | NA | NA | ornamental and horticultural plants | NA | NA | FALSE | FALSE | Lactuca sativa |

**Figure S3:** Example of information extracted from an abstract of a study by Landal et al. (2024), highlighting the extraction of the pathogenic vectors and the locations where a herbivorous pest is stated to be invasive. In addition to its role as a major pest of horticultural crops, its association with lettuce is correctly extracted (with regards to which the herbivore is not explicitly mentioned to be a pest). The nematode *Angiostrongylus costaricensis* is correctly inferred as a parasite.

NATURAL ENEMIES OF PENTALONIA NIGRONERVOSA COQUEREL, A VECTOR OF BUNCHY TOP OF BANANA AND BIOLOGY OF ITS MOST EFFECTIVE PREDATOR SCYMNUS NUBI LUS MULSANT. Abstract: Natural enemies of Pentalonia nigronervosa Coquerel (Hemiptera: Aphididae), a vector of bunchy top of banana, are reported from Tamil Nadu, South In dia. Totally four predators including three coccinellids (Pseudaspidimerus trinotatus (Thunberg), Scymnus (Scymnus) nubilus Mulsant, Cheilomenes sexmaculata (F.)) and one h emerobiid (Micromus timidus Hagen) were recorded. Only one parasitoid, Aphelinus sp. nr. gossypii Timberlake (Hymenoptera: Aphelinidae) was recorded. Biology of S. nubilus, the most effective predator of P. nigronervosa, was also studied in the laboratory and the results are presented. © 2023 Elsevier B.V., All rights reserved.. Author keywords: Aphid; Banana; Biology; Bunchy Top; India; Parasitoid; Population Dynamics; Predators; Predatory Potential; Scymnus Nubilus.

| Subject | Object | Type | Feeding Mode | Is Pest | Pest Importance | Herbivore Is BCA | Associated Industries | Invasive In | Vectors | Important Enemy | Biocontrol | Associated Plants |
|---|---|---|---|---|---|---|---|---|---|---|---|---|
| Pentalonia nigronervosa | Banana (Musa spp.) | herbivore | herbivory | TRUE | major | FALSE | fruit plantations | NA | Banana bunchy top virus (BBTV) – causing Bunchy Top disease of banana | NA | NA | NA |
| Pseudaspidimerus trinotatus | Pentalonia nigronervosa | natural enemy | predation | NA | NA | NA | fruit plantations | NA | NA | FALSE | FALSE | Banana |
| Scymnus nubilus | Pentalonia nigronervosa | natural enemy | predation | NA | NA | NA | fruit plantations | NA | NA | TRUE | FALSE | Banana |
| Cheilomenes sexmaculata | Pentalonia nigronervosa | natural enemy | predation | NA | NA | NA | fruit plantations | NA | NA | FALSE | FALSE | Banana |
| Micromus timidus | Pentalonia nigronervosa | natural enemy | predation | NA | NA | NA | fruit plantations | NA | NA | FALSE | FALSE | Banana |
| Aphelinus gossypii | Pentalonia nigronervosa | natural enemy | parasitism | NA | NA | NA | fruit plantations | NA | NA | FALSE | FALSE | Banana |

**Figure S4:** Example of information extracted from an abstract of a study by Poorani et al. (2023), highlighting the extraction of an important natural enemy (but not biocontrol agent) among multiple other natural enemies. *Scymnus nubilus* is correctly identified as an important natural enemy, as it was found in the study to be "the most effective predator of P. nignonervosa." Moreover, the herbivore *Pentalonia nigronervosa* was correctly identified as a vector of banana bunchy top virus. The extraction of *Pentalonia nigronervosa* as a major pest of banana is assumed to follow the prompt instructions that the subject of a biocontrol study may be inferred to be a pest. A search of the literature verifies that the herbivore is considered a major pest of bananas due to its vectoring of banana bunchy top virus (Mathers et al., 2020).

The potential of sympherobius pygmaeus (Rambur, 1842) as a biological agent against planococcus citri (Risso, 1813) in citrus orchards. Abstract: Planococcus citri (Risso) (He miptera: Pseudococcidae) is one of the most significant pests for especially citrus, crop plants and ornamental plants. Besides its worldwide importance, chemical control is a s ignificant, major method to suppress the population of P. citri in Turkey. Moreover, biological control has become more important in recent years because of the increase in cons umer consciousness about pesticides. For this reason, the population dynamic of P. citri and its predator, Sympherobius pygmaeus (Rambur) (Neuroptera: Hemerobiidae) in thre e different grapefruit orchards were studied. For the study, 10 trees were randomly chosen from each orchard and the individuals on the plants, 4 branches, and stem of each tre e were counted to determine the population level. When P. citri population reached the peak level, approximately 500 S. pygmaues individuals were released to each orchard. Po pulation levels of prey and predators were monitored for two years, with weekly intervals between May-October and one time for two weeks between December and April of the following year. As the result of the treatment process, it is observed that the predator S. pygmaeus succeeded in decreasing the population of P. citri under economic damage threshold level. As a result, it is demonstrated that the predator S. pygmaeus can be used as a biological agent against P. citri in integrated pest management programs as an environmentally friendly method. © 2020 Elsevier B.V., All rights reserved.. Author keywords: Biological Control; Grapefruit; Hemerobiidae; Mealybug; Neuroptera; Population.

| Subject | Object | Type | Feeding Mode | Is Pest | Pest Importance | Herbivore Is BCA | Associated Industries | Invasive In | Vectors | Important Enemy | Biocontrol | Associated Plants |
|---|---|---|---|---|---|---|---|---|---|---|---|---|
| Planococcus citri | Citrus paradisi (grapefruit) | herbivore | herbivory | TRUE | major | FALSE | fruit plantations, unspecified agriculture, ornamental and horticultural plants | NA | NA | NA | NA | NA |
| Planococcus citri | agricultural crops | herbivore | herbivory | TRUE | major | FALSE | fruit plantations, unspecified agriculture, ornamental and horticultural plants | NA | NA | NA | NA | NA |
| Planococcus citri | ornamental plants | herbivore | herbivory | TRUE | major | FALSE | fruit plantations, unspecified agriculture, ornamental and horticultural plants | NA | NA | NA | NA | NA |
| Sympherobius pygmaeus | Planococcus citri | natural enemy | predation | NA | NA | NA | fruit plantations | NA | NA | TRUE | TRUE | Grapefruit, citrus |

**Figure S5:** Example of information extracted from an abstract of a study by Yayla et al. (2020), highlighting the extraction of biocontrol provision. *Sympherobius pygmaeus* is correctly identified as a predator of the citrus pest *Planococcus citri* that was observed in the study to have "succeeded in decreasing the population of P . citri under economic damage threshold level", demonstrating that the enemy "can be used as a biological agent against P . citri in integrated pest management programs." *Planococcus citri* is correctly identified as a major pest of ornamental plants and (unspecified) agricultural crops. Whilst its association with grapefruit is apparent from the study area in grapefruit orchards, the herbivore's role as a major pest of citrus fruits in general was missed.

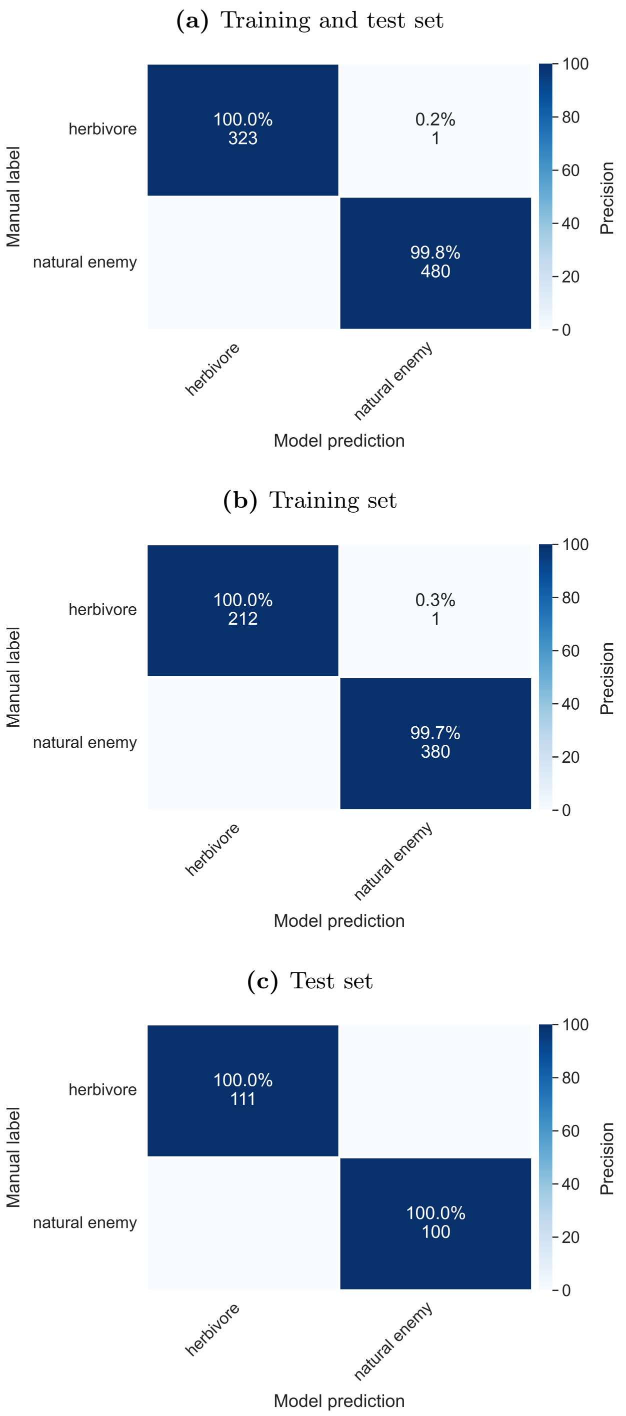


**Figure S6:** Confusion matrices of the column 'Species Type', computed on both the training and test set (a), the training set only (b) and the test set only (c). Percentages and colour scales refer to precision scores (%), which indicate what percentage of labels predicted (extracted) by the model were true positives, if a label was predicted by the model: Precision = TP/(TP+FP).

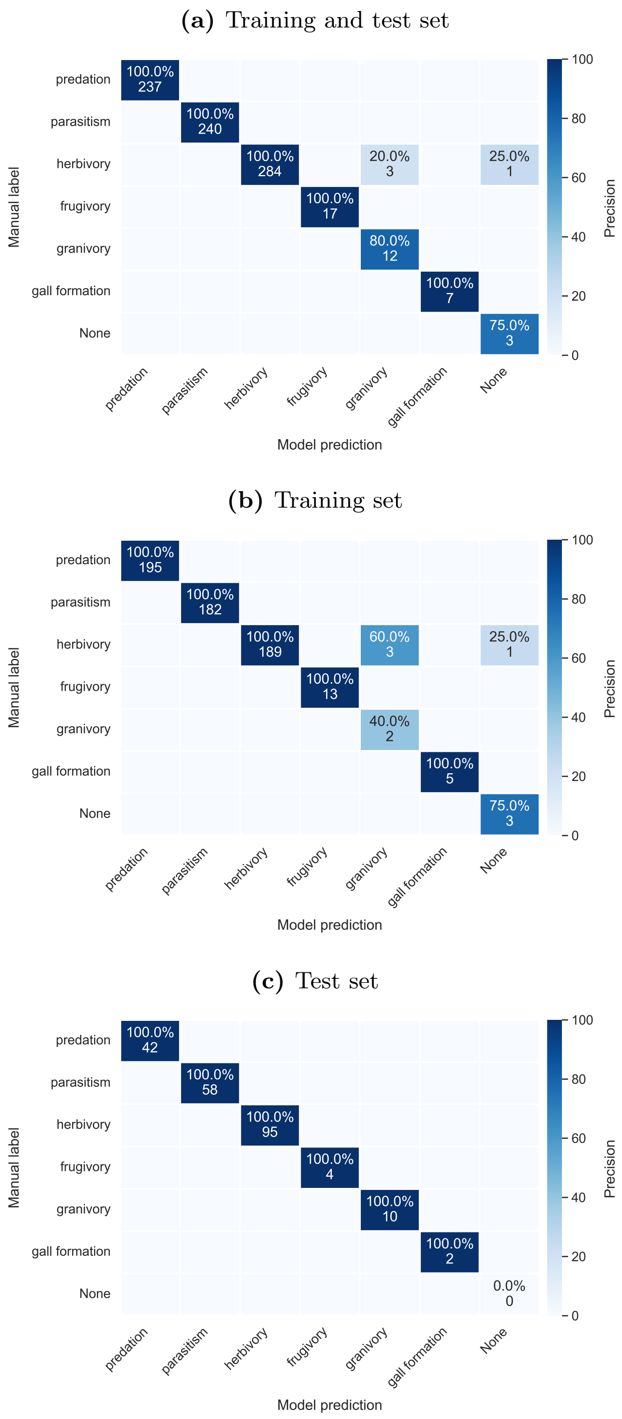


**Figure S7:** Confusion matrices of the column 'Feeding Mode', computed on both the training and test set (a), the training set only (b) and the test set only (c). Percentages and colour scales refer to precision scores (%), which indicate what percentage of labels predicted (extracted) by the model were true positives, if a label was predicted by the model: Precision = TP/(TP+FP).

**(a)** Training and test set

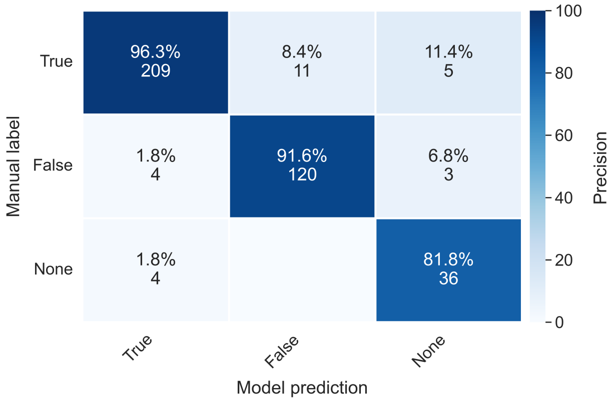


**(b)** Training set

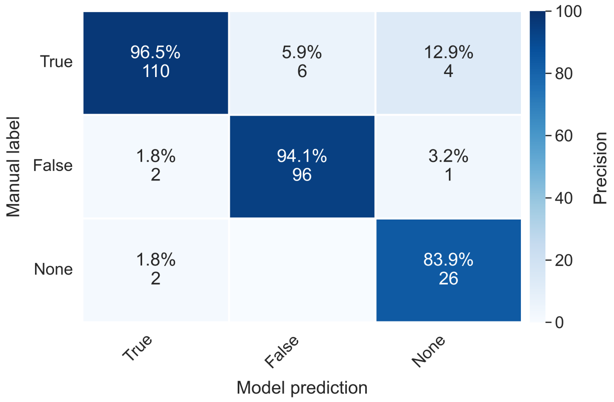


**(c)** Test set

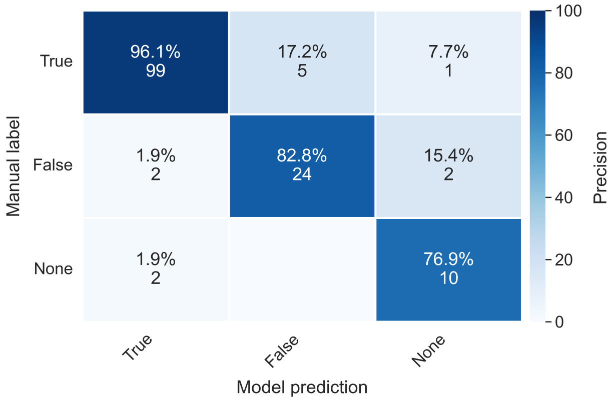


**Figure S8:** Confusion matrices of the herbivore column 'Is Pest', computed on both the training and test set (a), the training set only (b) and the test set only (c). Percentages and colour scales refer to precision scores (%), which indicate what percentage of labels predicted (extracted) by the model were true positives, if a label was predicted by the model: Precision = TP/(TP+FP).

**(a)** Training and test set

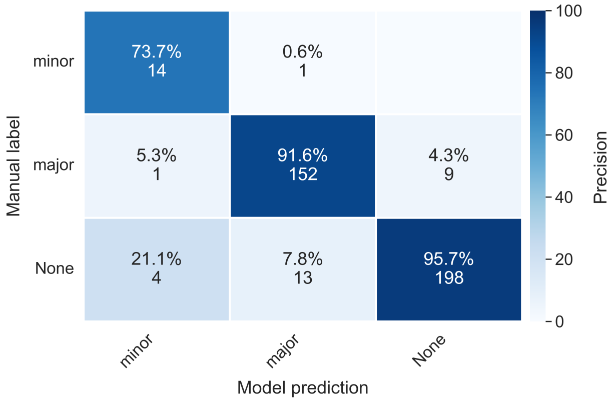


**(b)** Training set

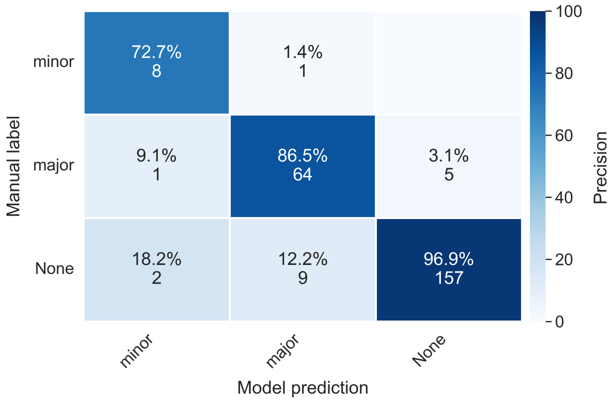


**(c)** Test set

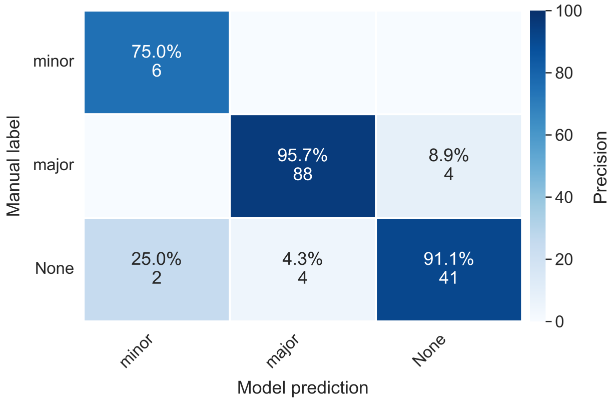


**Figure S9:** Confusion matrices of the herbivore column 'Pest Importance', computed on both the training and test set (a), the training set only (b) and the test set only (c). Percentages and colour scales refer to precision scores (%), which indicate what percentage of labels predicted (extracted) by the model were true positives, if a label was predicted by the model: Precision = TP/(TP+FP).

**(a)** Training and test set

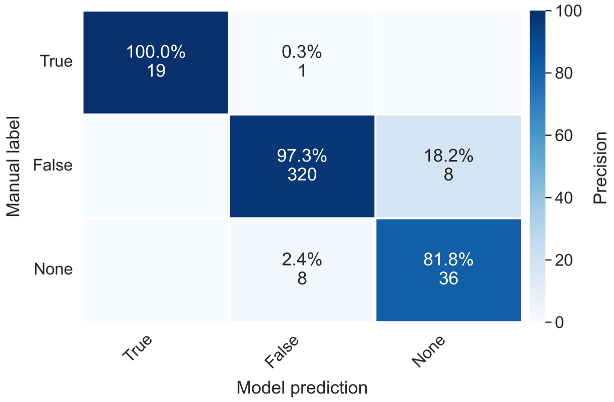


**(b)** Training set

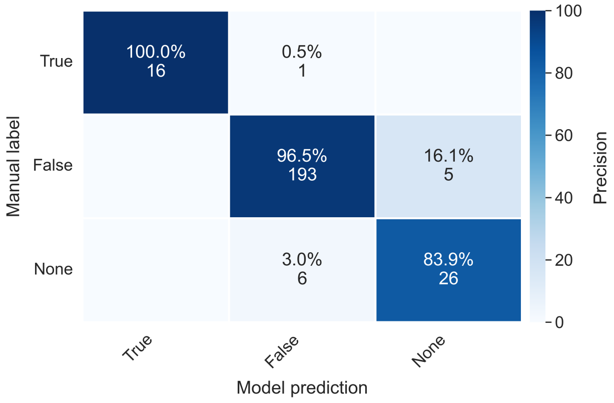


**(c)** Test set

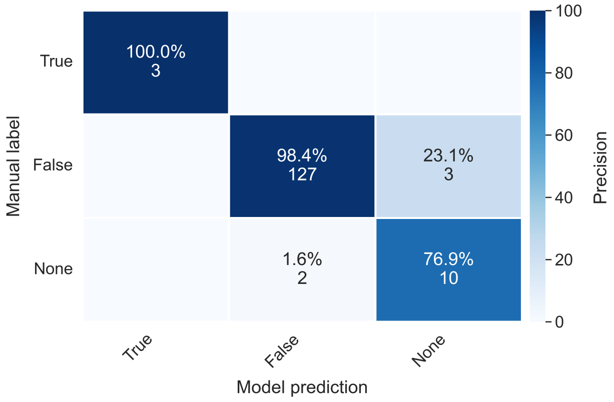


**Figure S10:** Confusion matrices of the herbivore column 'BCA (Biological Control Agent)', computed on both the training and test set (a), the training set only (b) and the test set only (c). Percentages and colour scales refer to precision scores (%), which indicate what percentage of labels predicted (extracted) by the model were true positives, if a label was predicted by the model: Precision = TP/(TP+FP).

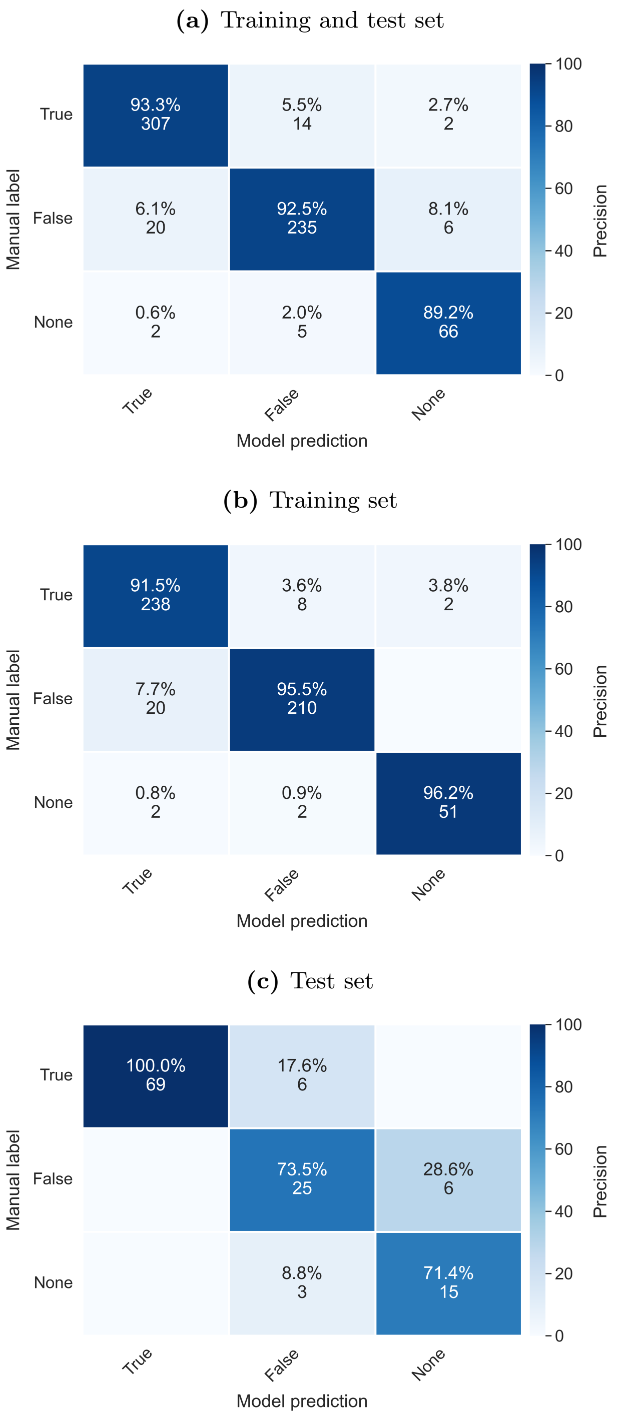


**Figure S11:** Confusion matrices of the natural-enemy column ‘Important Enemy’, computed on both the training and test set (a), the training set only (b) and the test set only (c). Percentages and colour scales refer to precision scores (%), which indicate what percentage of labels predicted (extracted) by the model were true positives, if a label was predicted by the model: Precision = TP/(TP+FP).

**(a)** Training and test set

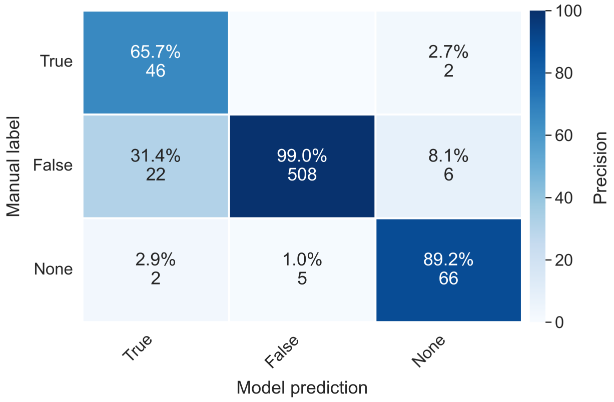


**(b)** Training set

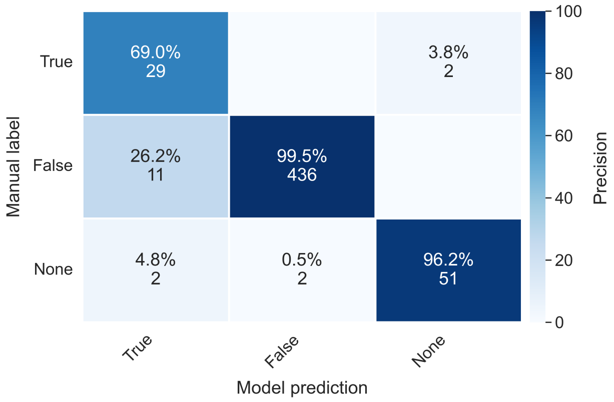


**(c)** Test set

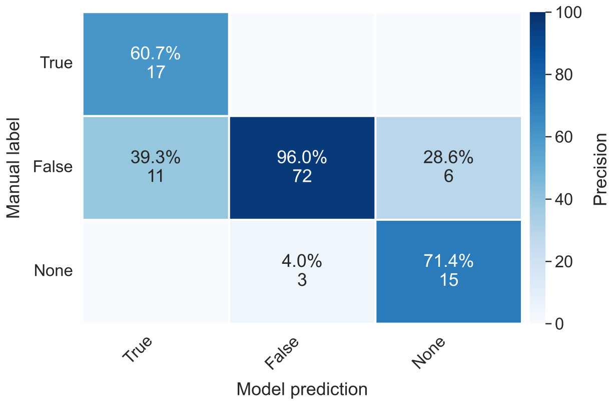


**Figure S12:** Confusion matrices of the natural-enemy column 'Biocontrol', computed on both the training and test set (a), the training set only (b) and the test set only (c). Percentages and colour scales refer to precision scores (%), which indicate what percentage of labels predicted (extracted) by the model were true positives, if a label was predicted by the model: Precision = TP/(TP+FP).

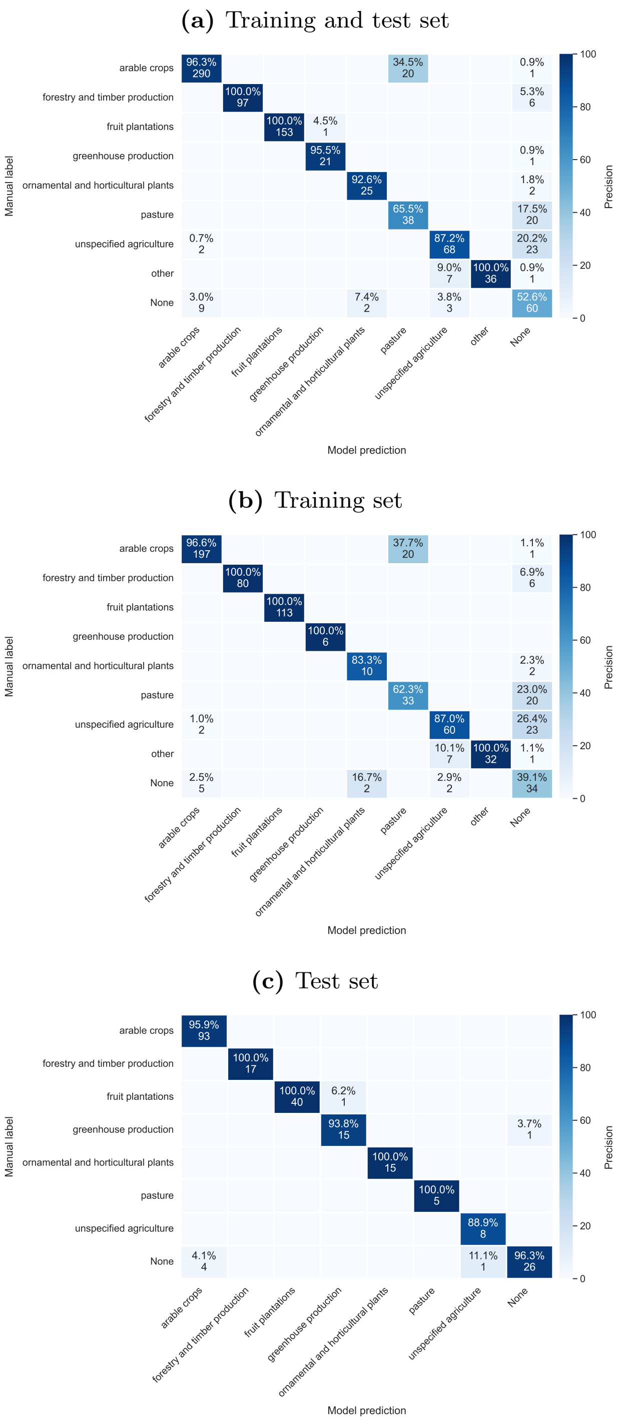


**Figure S13:** Confusion matrices of the column 'Associated Industries', computed on both the training and test set (a), the training set only (b) and the test set only (c). Percentages and colour scales refer to precision scores (%), which indicate what percentage of labels predicted (extracted) by the model were true positives, if a label was predicted by the model: Precision = TP/(TP+FP).
